# Mutation bias in driver genes reveals the distribution of effects of oncogenic mutations

**DOI:** 10.1101/2025.09.25.678277

**Authors:** Anastasia V. Stolyarova, Georgii A. Bazykin

## Abstract

Although many cancer driver genes have been identified, the full set of mutations within these genes that can promote tumorigenesis remains unknown. The contribution of a mutation depends on both its probability of occurrence and the selective advantage it confers. Here, we introduce a metric that quantifies how selection biases the spectrum of observed mutations in driver genes, and use it to develop a method estimating the number of driver mutations and their effect distribution at individual genes. Applying this framework to large cancer cohorts, we find that in most oncogenes, nearly all driver mutations have already been observed, whereas in most tumor-suppressor genes, the majority remain undiscovered. These results reveal fundamental differences in how mutation and selection shape mutational patterns across gene classes and provide a framework for interpreting newly detected variants.

## Main text

Tumor development is initiated when somatic cells acquire driver mutations that promote cell proliferation and therefore confer a selective advantage^1–3^. Identifying these mutations is critical to understanding of cancer biology and to diagnosis and treatment. However, up to half of all driver mutations remain undiscovered, and known drivers are often obscured by a larger number of neutral passenger mutations^4–6^.

The likelihood of observing a mutation in cancer is shaped by two forces: the rate at which the mutation arises and the strength of selection acting upon it^7–9^. Mutation rates vary substantially across nucleotide contexts due to differences in DNA damage and repair^10–13^. Selection within driver genes is also heterogeneous. Both genome-wide and experimental studies suggest that the distribution of fitness effects (DFE) varies widely across genes and tumor types^14–16^. Large-scale cancer sequencing has enabled development of statistical methods for identifying driver genes as well as specific mutations under positive selection while accounting for variation in local mutation rates^4,17–23^. These methods typically rely on patterns of recurrence, predicted functional impact, or deviations from neutral mutational expectations. However, they are primarily designed to identify the genes under selection rather than quantifying potentially advantageous mutations within a gene.

In this study, we aim to move beyond identifying driver genes to quantitatively characterizing their fitness landscapes. Specifically, we estimate two fundamental properties of the fitness landscapes of driver genes: the number of mutations that could act as drivers and the distribution of their fitness effects. Using the approximate Bayesian computation (ABC) framework, we infer gene-specific DFEs from the observed distributions of mutations. Our approach provides a quantitative summary of the selective landscape of driver genes, and enables predicting the number of yet-to-be-discovered driver mutations in these genes.

## Results

### Mutational spectrum bias in driver genes

To isolate the effect of selection on mutations in cancer genes, we first ask how patterns of mutations observed in driver genes differ from those in non-driver genes. We used TCGA, ICGC and PCAWG sequencing data to construct a dataset of missense somatic mutations in 13,510 samples of 25 tumor types (see Methods, Supplementary Fig. 1 and 2). For each tumor type, genes were classified as oncogenes (159 genes, 405 gene-tumor pairs), TSGs or ambiguous (243 genes, 754 gene-tumor pairs) or non-driver genes as predicted by the IntOGen pipeline^23^.

To estimate neutral mutational spectra of missense mutations, we used universal non-driver genes, *i*.*e*. those not annotated as drivers in any tumor type (18,472 genes), assuming that the effect of selection on distribution of somatic mutations within them is negligible (refs.^5,15^, but see ref.^24^). We estimated the probability of a missense mutation in each of the 96 three-nucleotide contexts by fitting a Poisson regression model (see Methods). We did this separately for each tumor type, thus controlling for the differences in mutation rates and spectra between tumor types^13,19^).

For each gene annotated as a driver in a tumor type and carrying at least 25 mutations in this tumor type, we then considered how the observed mutations were distributed across mutational contexts (Fig. 1a, green), asking if their distribution significantly differed from what was expected if they had accumulated according to their neutral mutation rates (Fig. 1a, gray). The strength of this deviation, which we refer to as mutational spectrum bias (MSB), in a gene is defined as the pseudo-R^2^, *i*.*e*. the additional variability explained by a model with fitted mutation rates as compared to the neutral null model (see Methods). This measure does not depend on the overall number of mutations observed, but instead asks whether these mutations have an unexpected distribution across contexts. As selection changes the distribution of mutations compared to the neutral expectation, we expect the strength of MSB to reflect the action of selection on mutations.

**Figure 1.**
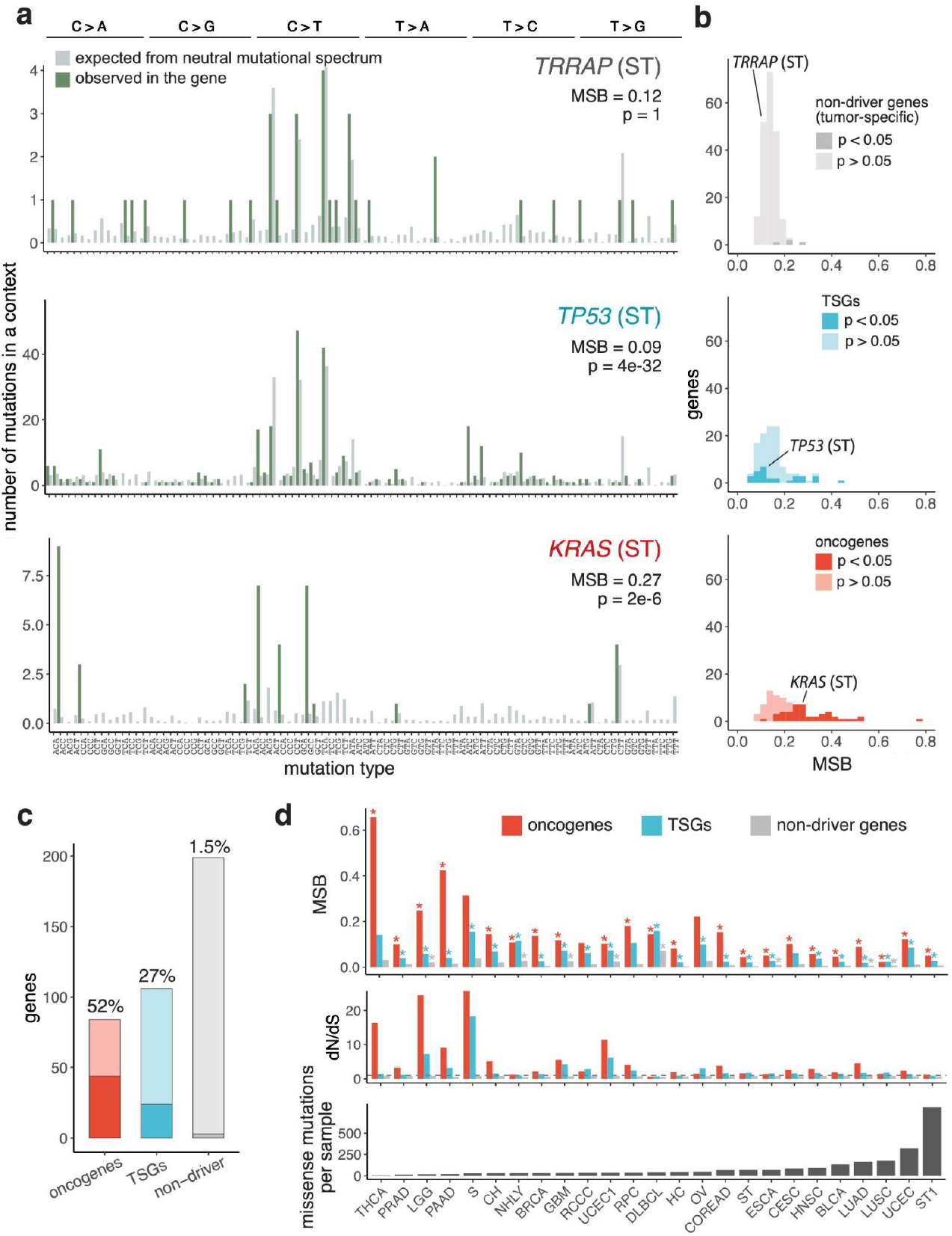
Mutational spectra in driver genes differ from the neutral spectra. **a**, Spectra of observed missense mutations in select genes across the trinucleotide contexts (green). The expected spectra are calculated from the genes that are not annotated as drivers in any tumor type and scaled to match the total number of observed mutations in the gene (gray). **b**, Strength of the mutational spectrum bias (MSB) in non-driver genes, TSGs and oncogenes. Only genes with 25 or more observed mutations within a tumor type are shown. **c**, Numbers of genes with significant MSB. For **b** and **c**, p-values were adjusted for multiple testing using Benjamini-Hochberg correction. **d**, MSB and dN/dS calculated in groups of genes pooled by tumor type and gene role. Asterisks show significantly biased spectra with p < 0.05.

The relative contribution of selection and mutation to the distribution of mutations varies across driver genes and is associated with the mechanism by which the gene affects tumor growth. Theory predicts that when the number of possible adaptive mutations (i.e., the target size) is small, selection will cause a stronger deviation from the neutral mutational spectrum, compared to the cases when this target is large^25–27^. Consistent with this prediction, tumor suppressor genes (TSGs), which have a broad range of potential loss-of-function mutations, are expected to have a weak MSB, *i*.*e*., a mutational spectrum of fixed mutations similar to the neutral spectrum, reflecting that the likelihood of observing a given mutation is primarily determined by the probability of this mutation to occur. By contrast, oncogenes typically have only a few sites that can acquire activating gain-of-function mutations. As a result, the distribution of mutations in oncogenes is shaped more by selective constraints than by mutation frequency, which is expected to result in a strong MSB^6^.

Indeed, the mutational spectrum bias is, on average, stronger in oncogenes than in TSGs (Figure 1b, signrank test p < 6e-10). Among the analyzed genes that are drivers in specific tumors, 52% (45/85) of oncogenes show a statistically significant deviation from the neutral mutational spectrum at a 5% FDR, compared to only 27% (30/114) of TSGs (Fig. 1c). As a control, we examined genes that act as drivers in other tumor types but not in the one analyzed (“tumor-specific non-drivers”) and found that only 1.5% (3/200) of these show significant MSB. Fig. 1a shows examples of mutational spectra in a non-driver gene with no detectable MSB (*TRRAP*), a moderately biased TSG (*TP53*) and a strongly biased oncogene (*KRAS*), all in stomach adenocarcinoma.

To compare the strength of MSB across tumor types, we grouped genes by functional role (oncogenes, TSGs, and non-drivers) and analyzed each tumor type separately. In nearly all tumor types, oncogenes show a stronger MSB than TSGs (Fig. 1d). By contrast, tumor-specific non-driver genes have either a weak (in 12 tumor types) or no (in 13 tumor types) bias at the 5% significance level. For both oncogenes and TSGs, the magnitude of the MSB is the highest in the cancer types with a low overall mutational burden (Spearman’s correlation p < 0.02). These tumor types also tend to have the highest dN/dS values in driver genes (Fig. 1d). This is consistent with previous observations that tumors with high mutational burdens typically carry only a small number of positively selected mutations, while most mutations are passengers^5,24^. As a result, the effects of selection on both dN/dS and MSB are obscured in such tumors.

### Mutational spectrum bias is driven by functional constraints

Another criterion used to identify genes under selection is mutation recurrence, the repeated observation of mutations at the same genomic site across tumor samples^14,18,28,29^. While recurrence of passenger mutations is well predicted by the background mutation rates, functionally important mutations driven by positive selection can be observed in multiple patient samples even if their expected mutation rates are low^5,15,19^. As with spectrum bias, recurrence patterns differ between gene classes: TSGs accumulate many rare and diverse mutations with little positional recurrence, while oncogenes typically carry a small number of highly recurrent activating mutations.

Consistent with the idea that recurrence and spectrum bias both reflect selection, we find that the strength of MSB is negatively correlated with the number of unique mutations observed per gene within both oncogenes and TSGs (Figure 2a; Spearman’s p = 1.6×10^-14^ for oncogenes, 4.8×10^-22^ for TSGs). Moreover, the spectrum bias is stronger in those genes where individual mutations recur more frequently across samples (Supplementary Fig. 3; *p* = 4.4×10^-4^ for oncogenes, 0.25 for TSGs). These results further support the view that strong spectrum bias reflects selection acting on a limited set of functionally advantageous sites (as also seen in simulations, Supplementary Fig. 4).

**Figure 2.**
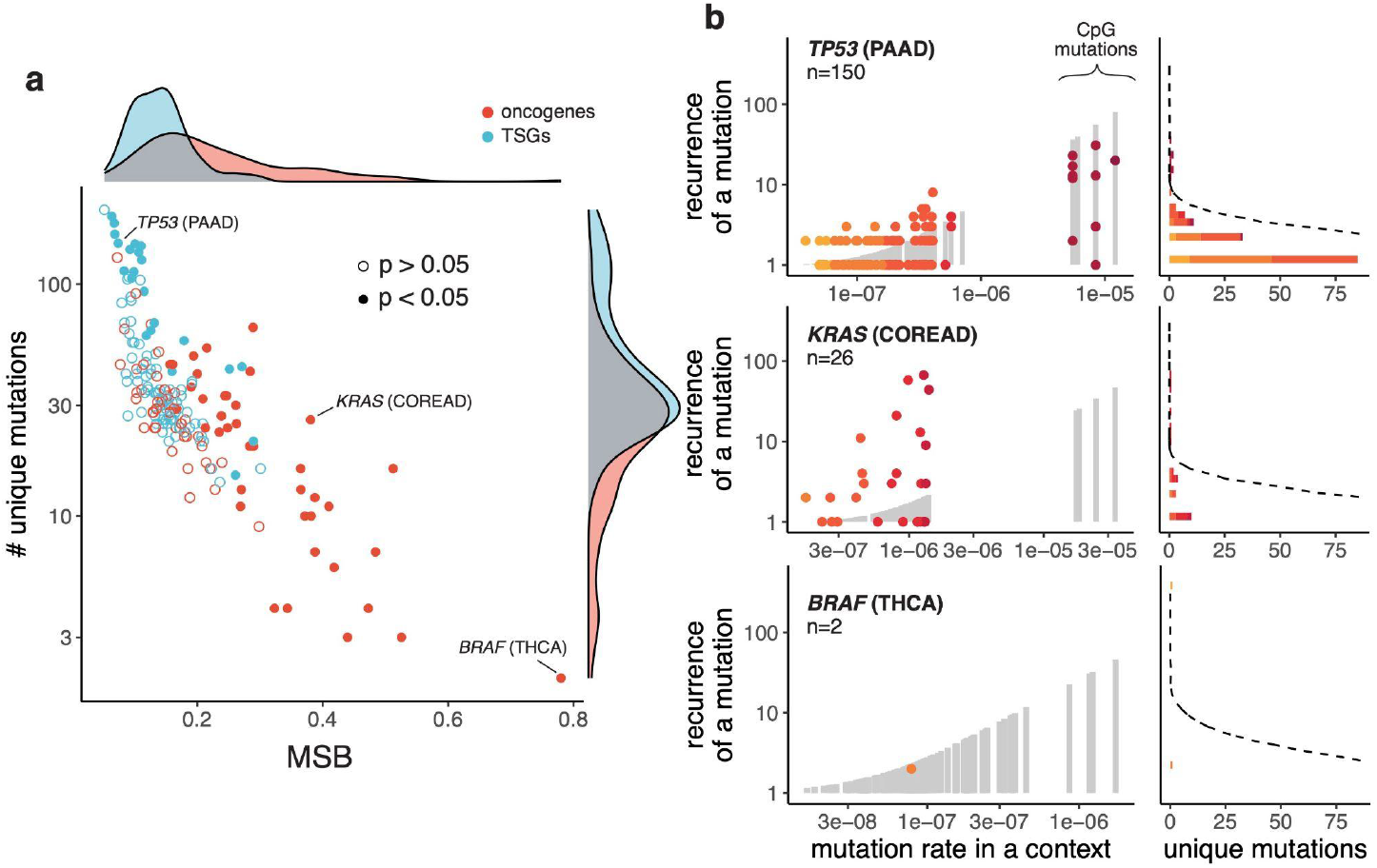
Mutational spectrum bias is stronger in driver genes with fewer unique mutations. **a**, The strength of MSB is negatively correlated with the number of unique mutations observed in a gene. Each point represents a gene in a tumor type; filled circles indicate genes with significant bias (FDR < 0.05). Only genes with at least 25 observed mutations are shown. **b**, Recurrence of observed mutations across sequence contexts in example genes with weak (TP53), moderate (KRAS) and strong (*BRAF*) spectrum bias. Each dot represents a unique mutation, with darker shades of red corresponding to higher-rate mutations. Grey columns and dashed lines indicate the expected distribution under the neutral mutational spectrum, matched for the total mutation count.

Fig. 2b shows examples of mutational patterns within individual driver genes. Each of these genes carries over a hundred mutations within a cancer type, but they differ in how these mutations are distributed across sites. In *BRAF*, where just two unique mutations are observed in the thyroid carcinoma (THCA) cohort, their occurrence is poorly explained by neutral mutational spectra: neither of them is a C>T transition in a CpG site, which have the highest mutation rate in this tumor type, and the more recurrent mutation is in the less mutable context. *KRAS* has more unique mutations, yet none of them fall within a CpG context, indicating that their high recurrence isn’t primarily caused by their high mutation rate. By contrast, in the *TP53* tumor suppressor gene, selection causes a high mutational burden but the distribution of mutations across sites is close to that expected under the neutral model, with the most recurrent mutations found in CpG contexts.

### Bias towards mutations with a low rate in driver genes

If selection and the mutational process act independently, every mutation that can occur within a driver gene is equally likely to be under positive selection regardless of its sequence context. If so, the probability that a mutation is functional does not depend on its rate of origin, and the expected distribution of positively selected mutations across sequence contexts should reflect the relative occurrence of these contexts in the gene.

However, highly mutable contexts are relatively rare in the genome. For example, C>T transitions in CpG sites account for between 7% and 48% of all missense mutations observed at non-driver genes depending on cancer type, but comprise only about 2% of all possible missense mutations in the genome (Fig. 3a). Therefore, if the set of possible driver mutations is small, most selected mutations will not fall into these high-mutability contexts, simply because such contexts are rare. As a result, the driver genes will, on average, appear enriched for mutations in low-mutability contexts, as confirmed by simulations (Supplementary Fig. 5).

**Figure 3.**
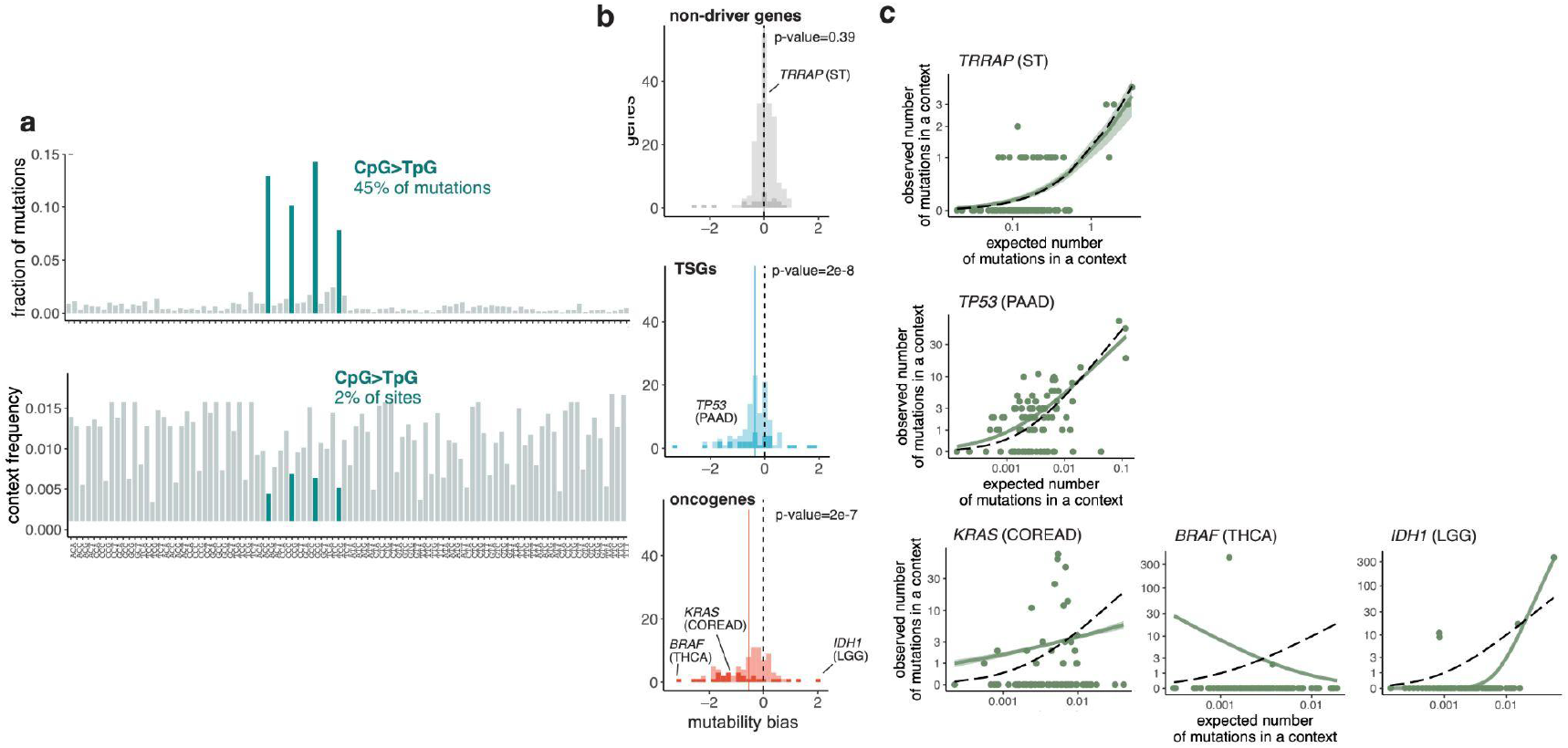
Mutations in driver genes are enriched in low-mutability sequence contexts. **a**, Frequencies of missense mutations across trinucleotide contexts in universal non-driver genes in pancreatic adenocarcinoma (PAAD) (top), compared to the frequency of corresponding contexts (bottom). **b**, The observed deviation from expected mutability in driver and non-driver genes with at least 25 mutations in the given tumor type. Mutability bias is calculated as the difference between the observed and expected average log mutation rate of the accumulated mutations. Negative values indicate that mutations preferentially accumulated in low-mutability contexts, and positive values, in high-mutability contexts. Vertical dashed lines show the null expectation (no shift), and solid lines show the mean of the observed distribution. P-values are calculated using the Wilcoxon signed-rank test. **c**, Examples of genes with no mutability bias (*TRRAP*), and genes with spectra enriched in low-mutability (*TP53, KRAS, BRAF*) or high-mutability (*IDH1*) contexts. Each dot represents a trinucleotide context. The dashed line shows the expected number of mutations under the neutral mutational spectrum (no mutability bias); the green line shows the Poisson regression fit to the observed data, with shaded areas showing standard error.

To test whether mutations in driver genes are systematically enriched in low- or high-mutability contexts compared to neutral expectations, we again used a Poisson regression model with a fitted mutability bias parameter defining the direction and strength of this shift (see Methods for details). Unlike the MSB which assesses any deviation from the neutral mutational spectrum, the mutability bias specifically measures whether the observed mutations preferentially occur in the sequence contexts with low or high expected mutation rates.

We find that in both TSGs and oncogenes, mutations preferentially occur in low-mutability contexts, while mutations in non-driver genes match the neutral expectation (Fig. 3b). 33% of oncogenes (28 genes), 26% of TSGs (30 genes), and 12% of non-drivers (23 genes) show a statistically significant enrichment in low or high-mutability contexts under the 5% FDR. In most of these cases, the shift is towards lower-mutability contexts (Fig. 3b-c). However, in some genes (5 oncogenes, 10 TSGs, and 8 non-drivers), mutations are significantly enriched in high-mutability contexts. For example, in *IDH1*, a highly recurrent mutation falls into a highly mutable CpG site (Fig. 3c). As predicted by simulations, this can occur when, in a minority of cases, positively selected sites happen to fall within highly mutable contexts, further amplifying their recurrence (Supplementary Fig. 5).

### Inferring the effects of mutations from mutational spectra

The variation in MSB and mutation recurrence across driver genes reflects how selection modulates the background mutational patterns. Although most individual sites do not accumulate enough mutations to estimate selection at single-mutation resolution, the MSB of a gene carries information about the fitness effects of the observed mutations. Here, rather than identifying specific driver mutations, we aim to reconstruct the distribution of fitness effects (DFE) for each gene – including both observed and unobserved mutations – capturing how many mutations are likely functional and how their effects vary in strength.

To infer gene-specific DFEs, we applied a simulation-based Approximate Bayesian Computation (ABC) approach. In this framework, the similarity between simulated and observed data is evaluated using a set of summary statistics: specifically, we used the total number of mutations in the gene, the number of unique mutations, their maximum recurrence, and the strength of MSB (see Methods for details). We described each gene’s DFE using three parameters: the number of mutations under positive selection *N*, the concentration parameter *α* (which is inversely related to the variance in fitness effects), and the scaling coefficient *s* which determines the overall strength of selection. To quantify the effective target size, we defined *N*^99^ as the number of distinct driver mutations expected to account for 99% of tumors with mutations in that gene. *N*^99^ captures both the number of positively selected sites and the variance of their fitness effects, jointly determined by *N* and *α* (see Methods).

Our method accurately infers *N*^99^ across simulated datasets (Pearson’s *R*^2^ between the true and estimated log-transformed values = 0.88; Supplementary Fig. 6), allowing us to estimate both the number and the distribution of driver mutations for each gene. Notably, omitting MSB from the model reduces inference accuracy, indicating that the spectrum bias captures information about selective constraints that is not reflected by recurrence patterns alone (Supplementary Fig. 6 vs. 7).

We applied ABC separately to each driver gene with at least 50 missense mutations in a given tumor type. On average, we infer that in oncogenes, just about 1% of all possible missense mutations in a gene account for 99% of tumors with mutations in that gene, reflecting strong selection acting on a very small number of sites (Fig. 4b,c). By contrast, for TSGs, about 9% of mutations contribute to 99% of cases, and their fitness effects tend to be more uniform.

**Figure 4.**
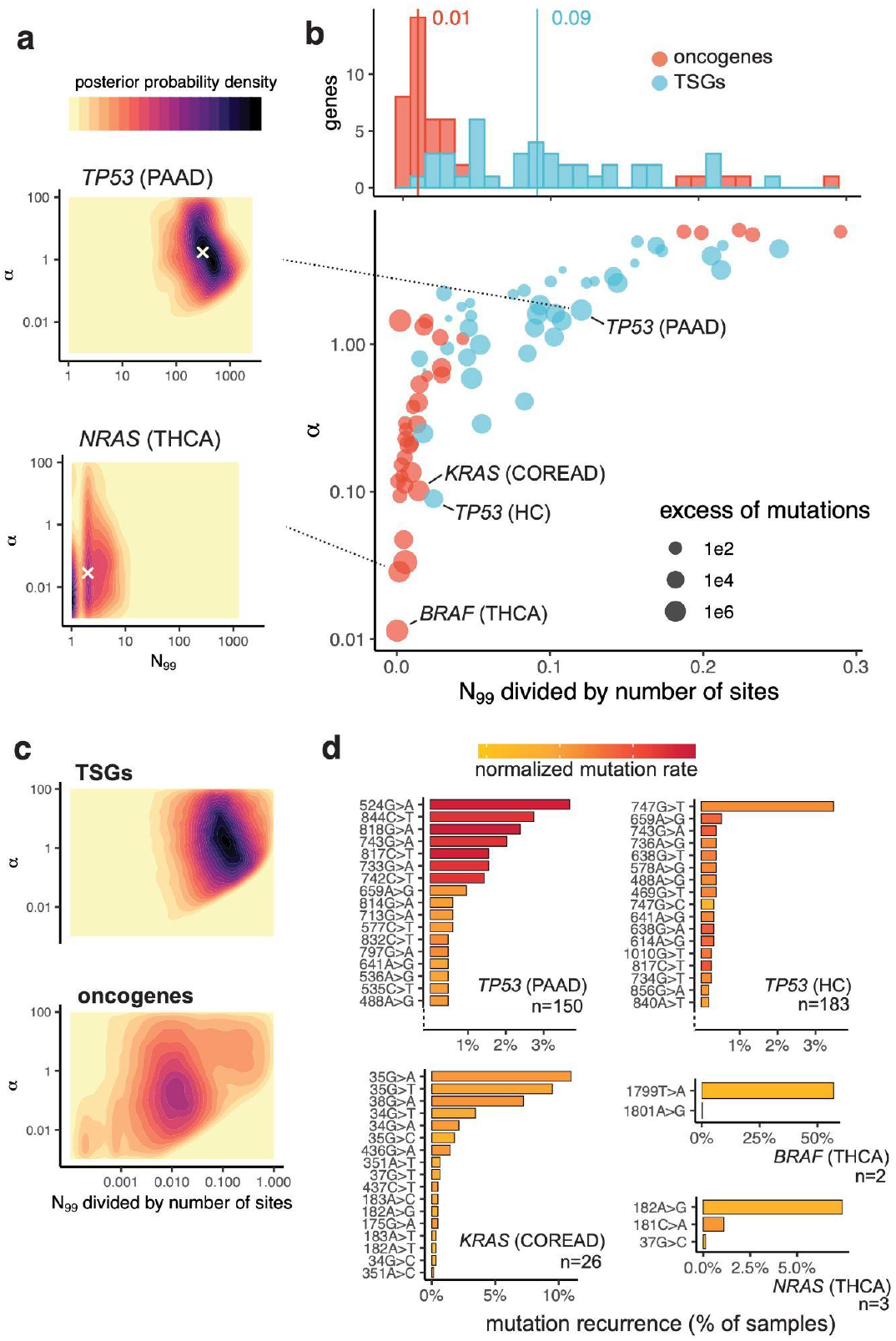
ABC estimates of the number of driver mutations and their effects across driver genes. **a**, Joint posterior distributions of the effective number of driver mutations *N*_*99*_ and concentration parameter □ estimated by ABC for a tumor suppressor gene *(TP53)* and an oncogene *(NRAS)*. Densities are based on 5,000 simulations accepted in the final ABC iteration. Distributions for all estimated 74 gene–tumor pairs with at least 50 mutations, and posterior distributions for *N* and scaling coefficient *s*, are shown in Supplementary Figs. 8-10. Both distributions and point estimates are available on https://github.com/astolyarova/drivers_abc. **b**, Estimated median values of *N*_*99*_ divided by the number of sites and of □ for the driver genes. Dot size indicates the fold increase in the number of missense mutations observed in the gene compared to that expected under neutral mutation rates. Vertical lines show the median fraction of *N*_*99*_ across TSGs and oncogenes. **c**, Pooled posterior distributions of *N*_*99*_ and *α* across genes, grouped by functional class. **d**, Mutations accumulated in genes with large *(TP53)*, intermediate *(KRAS)* and small *(BRAF, NRAS)* estimated *N*_*99*_. For *TP53*, only the 10 most recurrent mutations are shown.

These predictions align with the observed mutational patterns (Fig. 4d). Typical TSGs harbor a broad range of unique mutations with relatively uniform frequencies, and their most recurrent mutations tend to occur in highly mutable contexts – e.g., all top seven *TP53* mutations in pancreatic adenocarcinoma (PAAD) are CpG transitions. Oncogenes, by contrast, typically carry a small set of highly recurrent strong driver mutations, mostly outside high-mutability contexts, along with a tail of rarer, low-recurrence mutations. These rarer mutations are still unlikely to arise under neutrality, suggesting they too are under positive selection, albeit with weaker effects.

Notably, we did not identify any gene with only a few strong drivers and no tail of weakly selected ones (i.e., high *α* and small *N*^99^, falling in the upper left of Fig. 4b), even though our method can recover such cases (Supplementary Fig. 6). Even the most constrained oncogenes – such as *BRAF* and *NRAS* in thyroid carcinoma (THCA), where a single mutation accounts for over half of mutated samples – still carry additional low-frequency mutations likely representing weak drivers (Fig. 4d).

For genes annotated as drivers in multiple tumor types, we applied ABC independently for each tumor type. Despite the differences in neutral mutation rates and spectra across tumor types, the estimated parameters *α* and *N*^99^ for a given gene were consistent and showed no correlation with overall tumor mutability (Supplementary Fig. 11 and 12), supporting the robustness of our approach. An exception is *TP53*, which shows notably lower inferred *N*^99^ and *α* in hepatocellular carcinoma (HC) compared to other cancers such as pancreatic adenocarcinoma (PAAD), suggesting fewer driver sites with more variable fitness effects as typically observed in oncogenes (Fig. 4b). The most recurrent *TP53* mutation in HC (R249S, 747G>T) is strongly associated with aflatoxin B1 exposure^30^, which may alter the underlying mutational spectrum and bias our neutral model. Alternatively, our inference may reflect its known association with aggressive tumor progression and potential gain-of-function oncogenic activity of *TP53* in HC^31,32^.

### Predicting discovery rates of driver mutations

Previous studies suggest that up to half of all potential driver mutations remain undiscovered due to their low mutation rates or weak selective advantages^4,5^. As more tumors are sequenced, additional mutations continue to be found in known driver genes. However, not all of these are functional: distinguishing true drivers from passengers remains a central challenge, and several approaches have been developed to address it^5,14^. Meanwhile, the rate of discovery of new unique mutations has slowed with increased sample sizes, indicating that a large proportion of possible driver mutations may already have been observed^4,33^.

Using our estimates of the number and effect sizes of driver mutations in each gene, we can predict the rate at which new drivers will be uncovered as sample sizes increase (Fig. 5a; Supplementary Figs. 13 and 14). For genes with a large effective target size (*N*^99^) such as the tumor suppressor *TP53*, only a small fraction of all possible driver mutations have been observed to date. As a result, future sequencing is expected to reveal mostly new driver mutations. By contrast, for oncogenes with a small target size such as *BRAF, NRAS*, and *KRAS*, a large fraction of possible drivers has already been sampled. For these genes, further sequencing will yield diminishing returns, and most newly observed mutations will be passengers, particularly when the gene is under moderate selection (e.g., *FGFR3*). *N*^99^ estimates for all 74 gene–cancer pairs, together with the number of already sampled missense mutations, are shown in Fig. 6.

**Figure 5.**
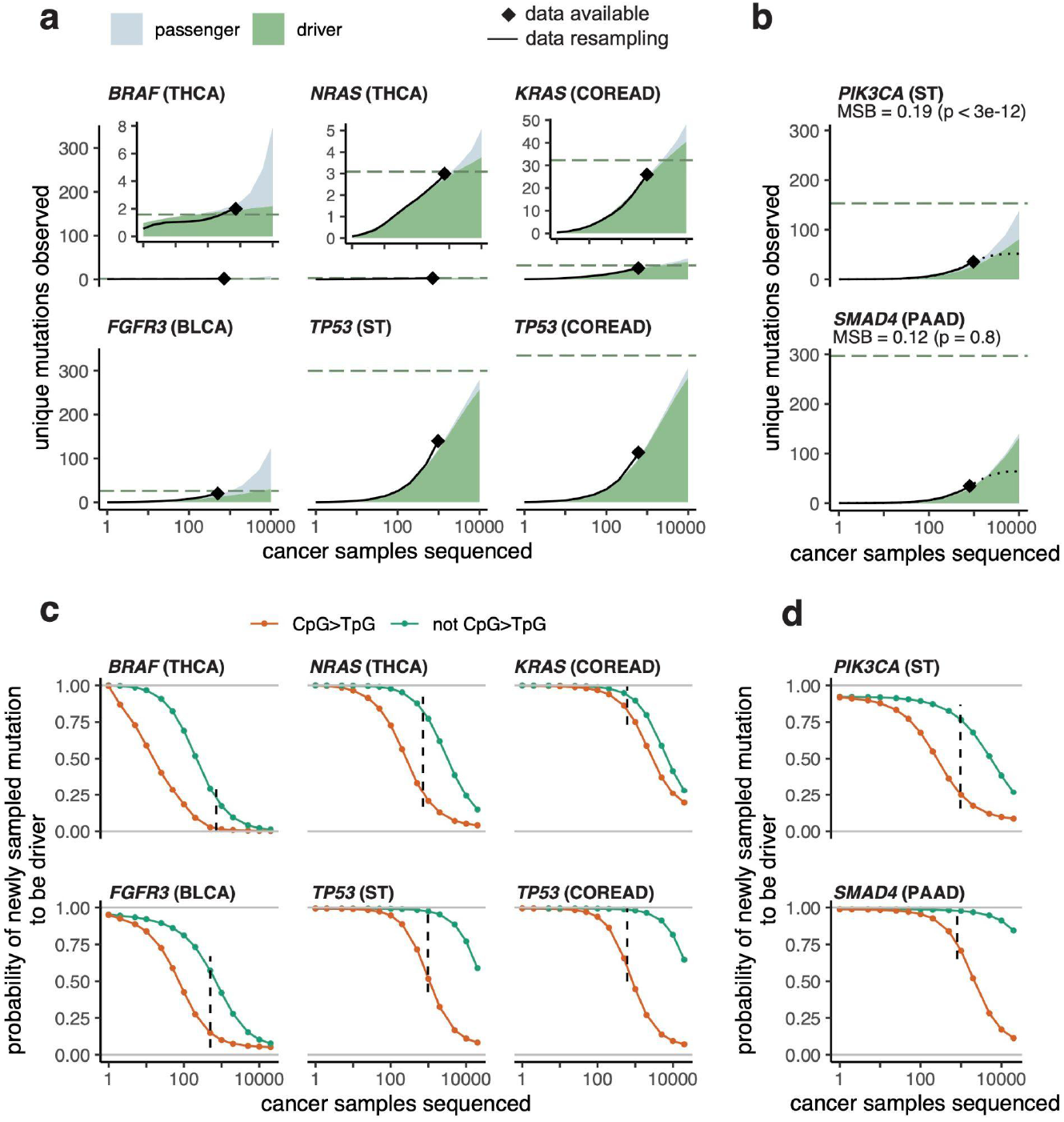
Predicting the number of driver mutations with increasing sample size. **a, b**, Observed and predicted numbers of unique mutations in selected driver genes. Diamonds indicate the total number of unique mutations observed in the analyzed dataset. Black lines show rarefaction curves from 1,000 bootstrap resamplings of tumor samples. Green areas indicate the number of driver mutations predicted to be found at this sample size; gray areas, passenger mutations. Predictions are based on 5,000 simulations using parameters from the final ABC iteration. Dashed green lines indicate the mean estimated *N*^99^. Predicted rarefaction curves for all genes are shown in Supplementary Figs. 13 and 14. **b**, Example genes with similar rarefaction curves but different MSB and therefore *N*^99^ values. Dotted black lines represent the saturation curves obtained by fitting an exponential function^6^. **c, d**, Predicted conditional probabilities that a newly sampled mutation is a driver, shown as a function of sample size and mutation context. Probabilities are estimated from simulations (see Methods). Vertical dashed lines indicate the currently available number of samples. Results for all driver genes are shown in Supplementary Figures 15 and 16.

**Figure 6.**
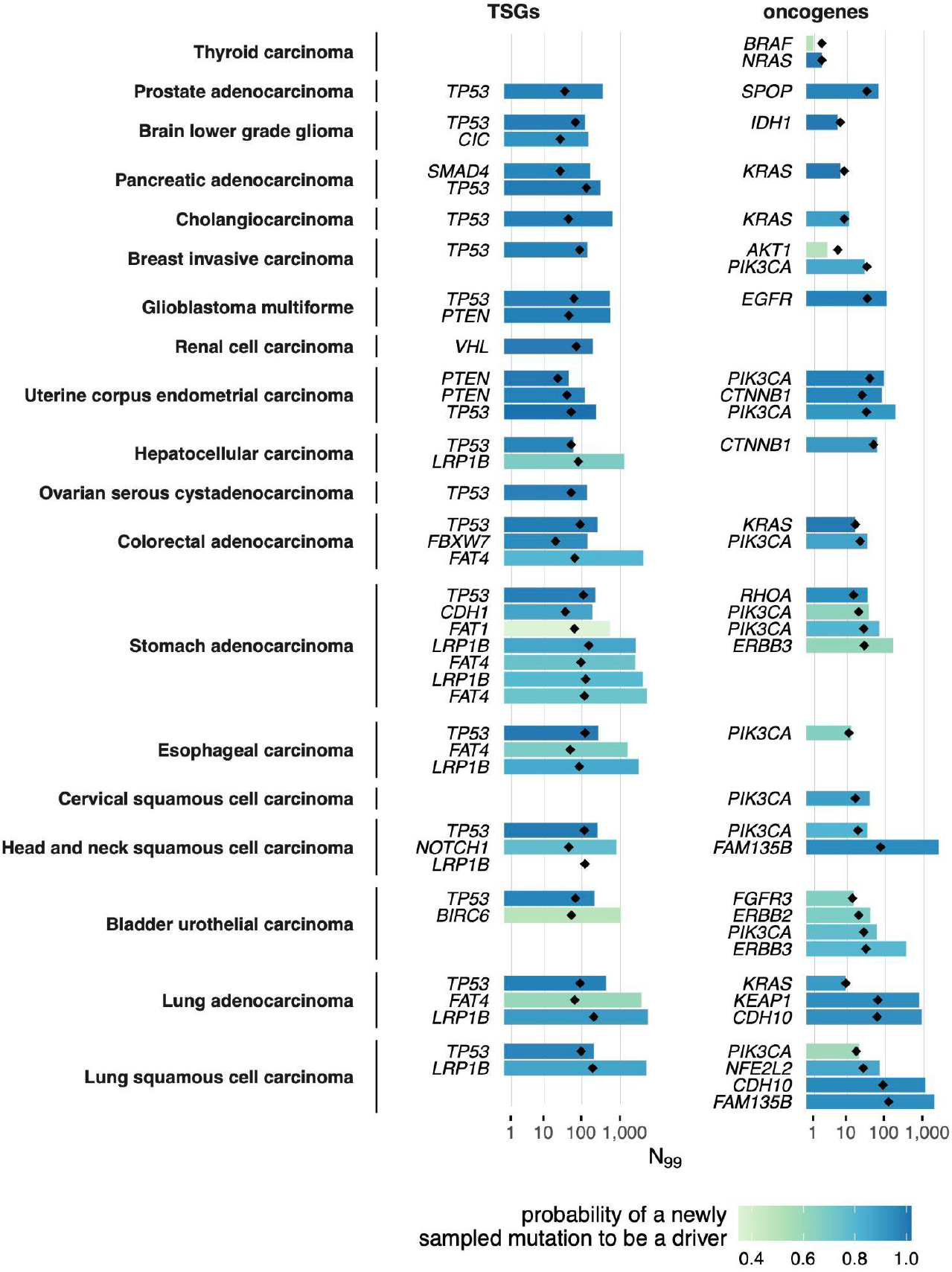
Predicted effective number of driver mutations in driver genes. For all 74 analyzed gene–tumor pairs, columns show the median estimate of the effective number of driver mutations *N*^99^. Diamond points show the number of unique missense mutations already sampled in the gene in the given cancer type dataset. The colors reflect the probability that a newly sampled mutation that wasn’t observed in the analyzed dataset before is a driver mutation (see examples in Figure 5c).

While saturation curves based on mutation accumulation rates can provide useful insights into the total number of driver mutations in a gene, they rely on mutation accumulation rates and may be misleading, particularly when the number of observed mutations is low. In these cases, incorporating MSB helps distinguish selective effects from neutral expectations. For example, the number of mutations observed in *PIK3CA* in stomach cancer and *SMAD4* in pancreatic adenocarcinoma increases at similar rates with sample size (Fig. 5b). However, the two genes differ markedly in their spectrum bias: *SMAD4*, as a typical tumor suppressor, shows no significant deviation from neutrality (p = 0.30), whereas *PIK3CA*, an oncogene, shows strong deviation (p = 3×10^-11^). Reflecting these differences, our method predicts that *SMAD4* has a much larger target size than *PIK3CA*. Consequently, at a sample size of 10,000 tumors, nearly all *SMAD4* mutations are expected to be drivers, while for *PIK3CA*, only about 50% are (Fig. 5b).

From the predicted saturation curves, we estimate the probability that a newly observed missense mutation in a gene is a driver (Fig. 5c,d, Supplementary Figures 15 and 16). This probability is initially high but declines as more tumors are sequenced, eventually reaching a point where most new mutations are expected to be passengers. The threshold at which this shift occurs is gene-specific, depending on the number and effect sizes of driver mutations. This probability also varies across mutation types: mutations in low-mutability contexts are unlikely to arise by chance and are more likely to be drivers, especially in small cohorts. By contrast, mutations in highly mutable contexts, such as transitions at CpG sites, are more frequently observed under neutrality and thus have lower posterior probabilities of being drivers.

The code for generating predictions of driver and passenger accumulation, as well as for computing driver probabilities by trinucleotide mutational context, is available on github (https://github.com/astolyarova/drivers_abc).

## Discussion

A change in the distribution of mutations compared to the background mutational process is a hallmark of selection. Models that incorporate non-uniformity of mutation rates, including biases in the rates of specific mutation types compared to their neutral expectations, have been previously used to identify cancer driver genes^13,19,22,23^. Here, we use the change in the distribution of mutations across sequence contexts to reconstruct the numbers and distributions of fitness effects of driver mutations in known driver genes.

Our inference is based on the interplay between selection and mutational constraints, reflected through deviations of mutations observed at driver genes from background mutational spectra. Perhaps unintuitively, these deviations are the strongest when the target for selection (i.e., the number of potential driver mutations) within the gene is small, because such selection focused on a few sites is unlikely to recapitulate existing mutational biases^25–27^. Furthermore, these deviations tend to be towards the mutations in low-mutability contexts, because the effect of selection inflating the number of observed mutations is stronger when the neutrally expected number of such mutations is low.

By combining recurrence data with mutational spectrum bias, we infer how many mutations in a gene are likely to be functional and how their selective effects vary. This perspective reveals systematic differences between gene classes: oncogenes typically experience strong selection acting on a few sites, while tumor suppressors are characterized by a wider set of weaker drivers. These differences align with the distinct functional roles of these gene groups in tumorigenesis.

Characterization of distributions of fitness effects of mutations within a gene facilitates the interpretation of these mutations, especially of those with low recurrence, underlying that functional importance is not captured by recurrence alone. Our results suggest that while a substantial fraction of driver mutations have already been sampled in well-characterized oncogenes, many remain undiscovered in tumor suppressors with larger target sizes. In these genes, newly sampled mutations, especially those in low-mutability contexts, should be candidates for functional follow-up.

## Methods

### Somatic mutation data

We collected somatic mutation data from tumor samples obtained by whole-exome (TCGA^34^ and ICGC^35^ datasets) or whole-genome (PCAWG^36^) sequencing. Samples were grouped into 46 tumor types following the IntOGen classification^23^, resulting in an initial dataset of 17,204 tumors from 122 patient cohorts. Only point mutations in coding sequences were included.

To minimize confounding by hypermutated samples, we excluded tumors with mutation counts exceeding 1.5 times the interquartile range above the third quartile for their respective tumor type, following ref.^23^. By this filtering, we excluded the 51 samples carrying known mutator variants in *POLE* gene, which drastically increase the mutation burden in the sample and affect the mutational spectrum^37,38^; *POLE* tumors comprised more than half of the excluded UCEC samples. Additionally, we removed duplicated samples. After this filtering, some tumor types demonstrated a bimodal distribution of the number of mutations per sample (Supplementary Fig. 1), indicating heterogeneity in mutational processes. For CH, PRAD, UCS, and S tumor types, we excluded samples from the hypermutated peak. For RCCC, ST, THCA, and UCEC, we split datasets into low- and high-mutation burden subsets. Tumor types with fewer than 50 samples or fewer than 5,000 total mutations were excluded from downstream analysis. Additionally, we excluded MEL (melanoma) due to its high inter-sample variability in mutational spectra and a large fraction of hypermutated samples^10^ (Supplementary Fig. 2). ACC and AML were excluded due to low mutation counts per sample (Supplementary Fig. 1).

The final dataset consisted of 13,510 samples grouped into 25 tumor types. All analyses were conducted separately for each tumor type.

Tumor-specific driver gene annotations were downloaded from the IntOGen database (release 2023.05.31)^23^. According to this annotation, driver genes are classified by their role as either oncogene or tumor suppressor gene (TSG). Genes labeled as “ambiguous” were pooled with the TSGs.

### Neutral mutational spectra

To estimate neutral mutational spectra, we selected the set of universal non-driver genes, *i*.*e*. the ones not annotated as drivers in any tumor type, assuming that the distribution of mutations within these genes is unaffected by selection and thus reflects neutral mutational processes.

For each of the 96 trinucleotide mutation types *c*, we estimated the neutral mutation rate *μ*_*c*_ using a Poisson regression model:

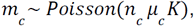

where *m*_*c*_is the total number of missense mutations of type *c* observed in universal non-drivers across all samples of the corresponding tumor type, *n*_*c*_ is the number of missense sites in the corresponding contexts, and *K* is the number of samples. Models were fitted independently for each tumor type.

Our model assumes that the neutral mutational spectrum and mutation rates are uniform across the genome, that is, the same for all genes within a given cohort exposed to the same mutational processes. While genes may differ in mutation rates due to local variation in mutational processes (e.g., replication timing, methylation level, etc.^10,13^), the mutational spectrum bias (MSB) is estimated solely from differences in spectra, and we do not expect such heterogeneity to affect the estimate. While local factors influencing the spectrum could, in principle, lead to overestimation of MSB and thus underestimation of the number of driver mutations in a gene (see below), we expect this effect to be small.

This model also assumes that all samples within a tumor type have the same mutational spectrum. To account for variation in mutational burden across samples, we implemented a heterogeneous model in which the per-sample mutation rate in each context scales with the overall sample mutability 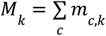 (*i*.*e*., the total number of missense mutations in non-driver genes in sample *k*):

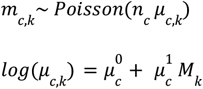

Here, *m*_*c,k*_ is the number of mutations of type *c* in sample *k*, and *μ*_*c,k*_ is the corresponding sample-specific mutation rate defined by intercept 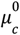 and slope 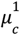.

For all tumor types except the ST dataset, the heterogeneous model significantly improved fit over the homogeneous model when applied to pooled data from universal non-driver genes (likelihood-ratio test p < 4×10^-4^). At the level of individual genes, however, neither model outperformed the other (p > 0.057), and the choice of model did not alter which driver genes were identified as having significantly biased spectra. Nevertheless, we used the heterogeneous model to estimate the MSB and compute associated p-values in all downstream analyses.

### MSB test

To assess whether a gene’s observed mutation spectrum deviates from neutral expectation, we compared two nested Poisson regression models using likelihood-ratio test (LRT). The null model assumes that positive selection increases the overall mutation rate uniformly across contexts without altering the spectrum:

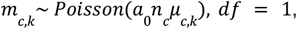

where *m*_*c,k*_ is the number of mutations of a given type in sample, *n*_*c*_is the number of sites, *μ*_*c,k*_ is the neutral rate estimated from universal non-driver genes, and *a*_0_ is a scaling factor reflecting the strength of selection.

The alternative model allows the spectrum of mutations observed in a gene to differ from the neutral spectrum by introducing context-specific scaling parameter *a*_*c*_:

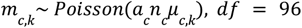

The significance of the MSB is estimated by comparison of models fit using LRT. The effect size of this bias is measured as McFadden’s pseudo-R^2^, indicating the improvement of the fitted model over the null model.

### Mutability bias

While the MSB test detects whether the observed mutation spectrum of a gene deviates from neutral expectations, it does not indicate the direction of this deviation. Specifically, we sought to determine whether mutations tend to occur more frequently in low-mutability or high-mutability contexts than expected under neutrality.

To do this, we fit an additional model in which context-specific mutation rates can be rescaled:

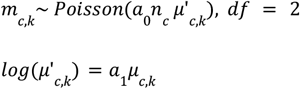

Here, *a*_1_ < 1 indicates enrichment in low-mutability contexts, while *a*_1_ > 1 indicates enrichment in high-mutability contexts. Genes for which this model provides a significantly better fit than the null model (based on the LRT) were classified as showing a mutability bias.

To quantify the strength of the mutability bias, we compare the average log mutation rate of observed mutations to that expected under neutrality, weighted by context frequency:

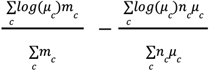

Negative values indicate enrichment in low-mutability contexts, whereas positive values indicate enrichment in high-mutability contexts.

### Simulations

We simulated accumulation of missense mutations within a gene, combining a neutral mutational spectrum with a gene-specific distribution of fitness effects (DFE). Each simulation assumed two categories of mutations: *N* driver mutations under positive selection, and all remaining mutations treated as neutral (passengers). Both driver and passenger mutations were randomly assigned to one of the 96 trinucleotide contexts according to their frequency in the gene’s coding sequence.

For each mutation *i* in sample *k*, its probability is defined as:

- for neutral mutations: *p*_*i,k*_ = *μ*_*c,k*_
- for selected mutations: *p*_*i,k*_ = *min*(*μ*_*c,k*_ *x*_*i*_ *s*, 1)

Here, *μ*_*c,k*_ is the context-specific neutral mutation rate in sample *k* (as estimated using the heterogeneous background model); *x*_*i*_ is the selection coefficient of mutation *i*; and *s* is a scaling factor capturing the overall strength of selection on the gene. Together, *x*_*i*_ *s* is the factor comparing the probability of occurrence of a positively selected mutation to a neutral mutation.

Selection coefficients *x*_*1*_, *…, x*_*N*_ were sampled from a Dirichlet distribution with concentration *N* parameters set to *α*, such that 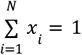. Thus, *α* defines the heterogeneity in fitness effects of driver mutations, with lower values of *α* corresponding to higher variance and therefore a skewed DFE, while *s* determines the cumulative strength of selection.

Our simulation model assumed that mutations are independent (*i*.*e*., cannot occur at the same site and do not interfere within a sample) and that selective pressure and mutation spectrum are constant over time.

The code for simulation is available at https://github.com/astolyarova/drivers_abc. Results illustrating the MSB as a function of *N* and *α* are shown in Supplementary Figs. 3 and 4.

### Estimation of DFE in driver genes

To infer the distribution of fitness effects (DFE) in driver genes, we employed an approximate Bayesian computation (ABC) framework. Inference was performed independently for each gene–tumor type pair with at least 50 observed missense mutations. Simulations incorporated the tumor-type-specific neutral mutational spectrum *μ*_*c,k*_, the trinucleotide context composition of the gene *n*_*c*_, and the distribution of mutational burden across samples *M*_*k*_ to capture heterogeneity in background mutability.

We jointly estimated three parameters: the number of positively selected mutations *N*, the concentration parameter of the Dirichlet distribution *α* controlling the variance of fitness effects, and the gene-level selection scaling coefficient *s*. Prior distributions were log-uniform: *N*ϵ [1, # *missense sites in the gene*], α ϵ [0. 01, 10], *s* ϵ [1*e*3, 1*e*9].

We used four summary statistics to compare observed and simulated data: (1) the total number of mutations, (2) the number of unique mutations, (3) the maximum recurrence of a mutation, and (4) the negative log-likelihood of the observed mutation spectrum under the neutral model. All summary statistics were log-transformed.

Parameter inference was conducted using the simulated annealing ABC (SABC) algorithm implemented in the ABCpy package^39,40^. ABC included up to 20 annealing steps or was halted if the acceptance rate fell below 1%. Each iteration produced 5,000 posterior samples. Default ABCpy settings were used unless specified otherwise. Posterior distributions of the estimated parameters were comprised of the 5,000 simulations from the final SABC iteration. The code is available at https://github.com/astolyarova/drivers_abc.

### Effective target size

The parameters *N* and □ capture different aspects of the DFE but both affect the size of the target for positive selection. When □ is small (*i*.*e*., the DFE is highly skewed), selection only favors a few strongly beneficial mutations even if *N* is large. To summarize the strength of selection constraint acting on a gene, we define the effective number of driver mutations, *N*_*99*_, as the number of driver mutations that together account for 99% of the cumulative selection:

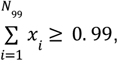

where *x*_*i*_ are normalized selection coefficients. When effects are evenly distributed (large *□*), *N*_*99*_≈N; when effects are heterogeneous (small *□*), *N*_*99*_≪N.

### Cross-validation of ABC

We validated our inference framework by simulating driver genes under known selection parameters and then re-estimating them using ABC. In these tests, the estimates of *N* and *-* were strongly correlated within datasets (Supplementary Fig. 8), reflecting the difficulty in disentangling a few evenly strong driver mutations from a broader set of driver mutations with skewed DFE.

By contrast, effective target size *N*_*99*_ was robustly inferred across a wide range of parameter combinations in simulation tests (Supplementary Fig. 6), showed narrower posterior intervals than *N* or □, and was largely independent of □ across both simulated and real data (Supplementary Figs. 6, 9, and 17). This makes *N*_*99*_ a more stable and interpretable summary of the gene’s effective selection target.

To assess the contribution of MSB to inference accuracy, we repeated the cross-validation excluding the spectrum likelihood from the summary statistics. This reduced accuracy in estimating both *N*_*99*_ and □ (Supplementary Figs. 6 and 7), indicating that spectrum bias captures additional information about the DFE in a gene.

### Discovery rate of new driver mutations

Using our estimates of the number and effect sizes of driver mutations in each gene, we predicted how the number of observed driver and passenger mutations scales with increasing sample size. For each gene, we simulated mutation accumulation across a range of sample sizes using posterior parameter distributions obtained from the ABC inference. Since each mutation in the simulation has a known selection coefficient, we tracked the number of unique driver and passenger mutations separately across simulated sample sizes (Figs. 5a-b; Supplementary Figs. 13 and 14).

Additionally, based on the estimated DFE in the gene, we calculated the probability that a new mutation is under positive selection given that it is observed in the *k*-th sample and has not been sampled before. This conditional probability is given by:

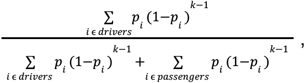

Where *p*_*i*_ is the probability for mutation *i* to be observed in the *k*-th sample, and 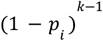 is the cumulative probability of it not being observed in the first (k − 1) samples. These probabilities were averaged across all posterior simulations to estimate the likelihood of discovering new drivers at each sample size.

This analysis was performed separately for different mutation classes. Discovery probabilities for CpG transitions versus all other mutation types are shown in Figure 5c-d and Supplementary Figs. 15 and 16. The code for computing discovery probabilities across trinucleotide contexts and sample sizes for each analyzed gene-tumor pair is available at https://github.com/astolyarova/drivers_abc.

## Acknowledgements

The results shown here are based on data generated by the TCGA Research Network, the International Cancer Genome Consortium and the Pan-Cancer Analysis of Whole Genomes projects. We thank Donate Weghorn and Ruslan Soldatov for comments on a draft of the manuscript.

