## Supplementary Figures for "Mutation bias in driver genes reveals the distribution of effects of oncogenic mutations"

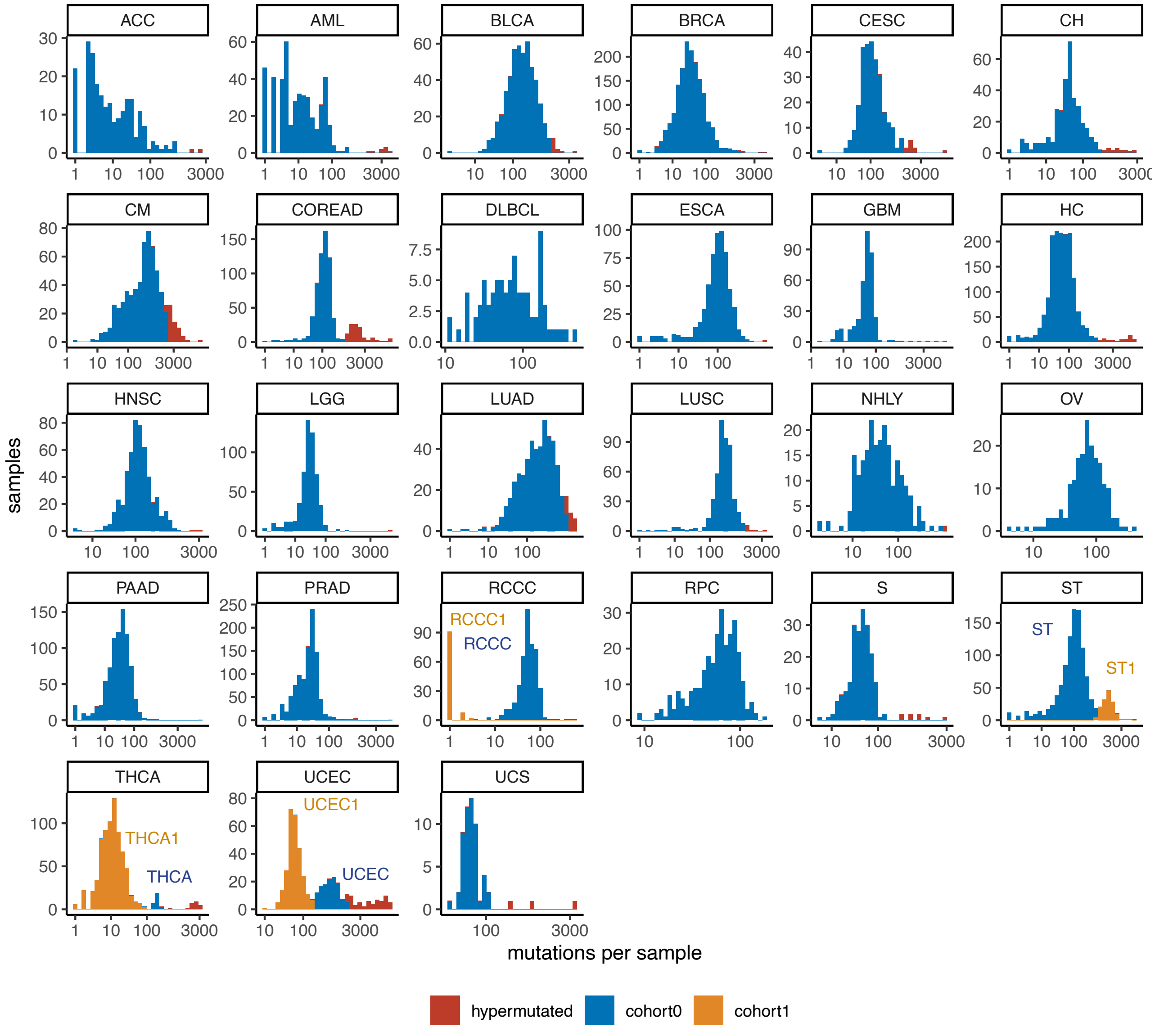


**Supplementary Figure 1. Mutational burden across analyzed tumor samples.** Only single-base substitutions (SBSs) in coding regions were included in the analysis. Tumor types with fewer than 50 samples or fewer than 5,000 total coding mutations are not shown. Samples with exceptionally high mutation counts (shown in red) are classified as hypermutated and excluded from further analysis. For ST, THCA, and UCEC, which exhibit a bimodal distribution in mutation burden, samples are split into two subsets, shown in blue and orange.

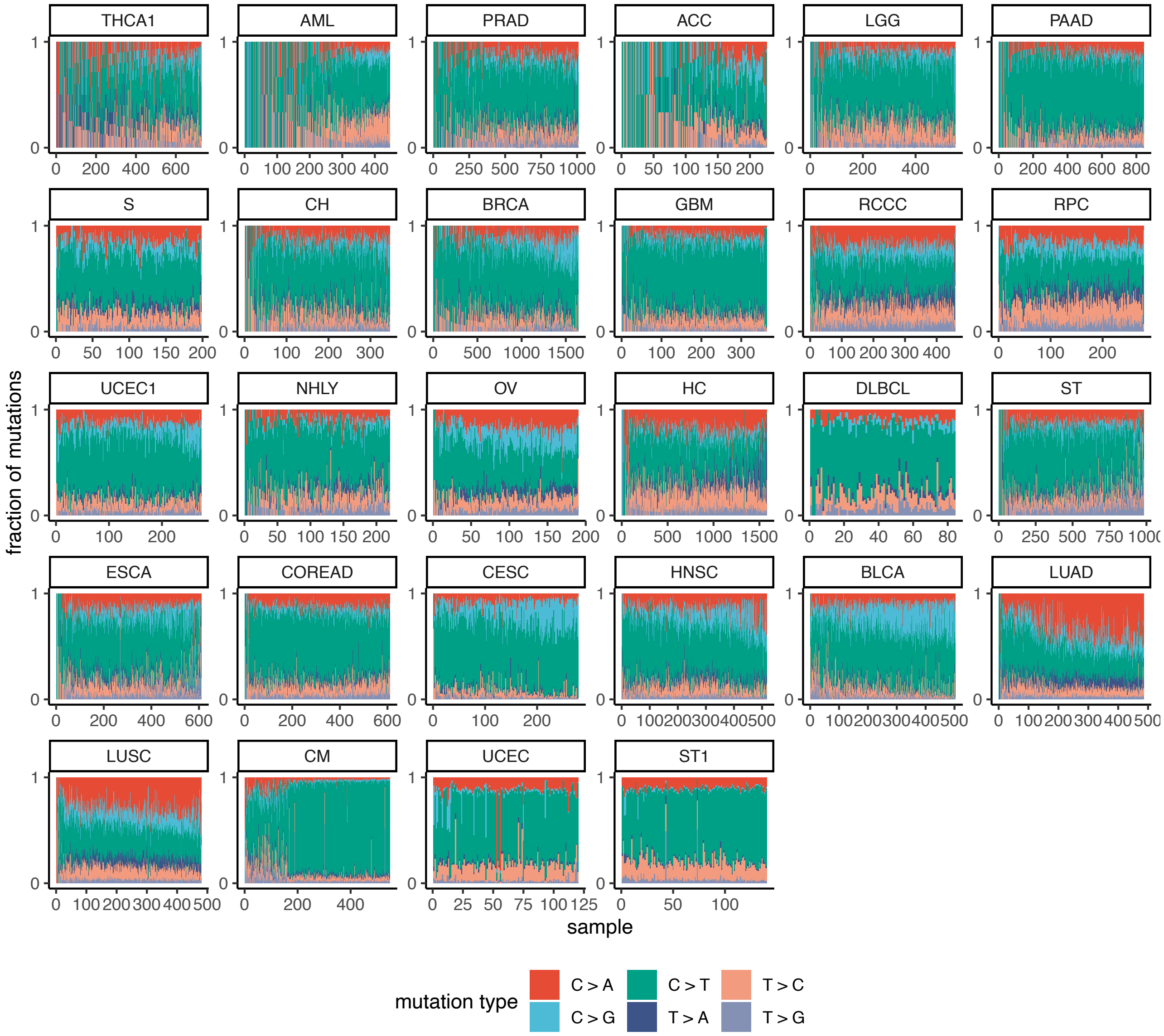


**Supplementary Figure 2. Mutational spectra across analyzed cohorts.**
Tumor types and individual samples within each cohort are ordered by average mutational burden per sample. Only single-base substitutions (SBSs) in coding sequences are included.

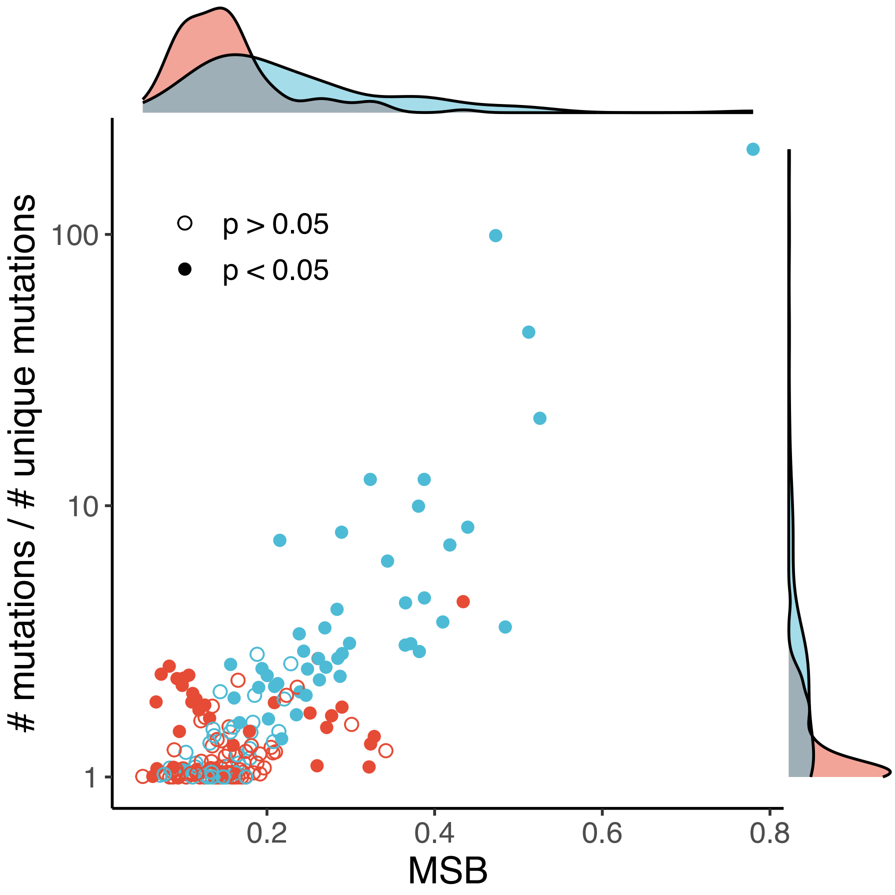


**Supplementary Figure 3. Mutational spectrum bias is stronger in driver genes with few highly recurrent mutations.**

Each point represents a gene in a given tumor type; filled circles indicate genes with significant mutational spectrum bias (likelihood-ratio rest $p < 0.05$ after multiple testing correction). Only genes with at least 25 observed mutations in a given tumor type are shown. The bias tends to be stronger in genes dominated by a small number of highly recurrent mutations (top right corner).

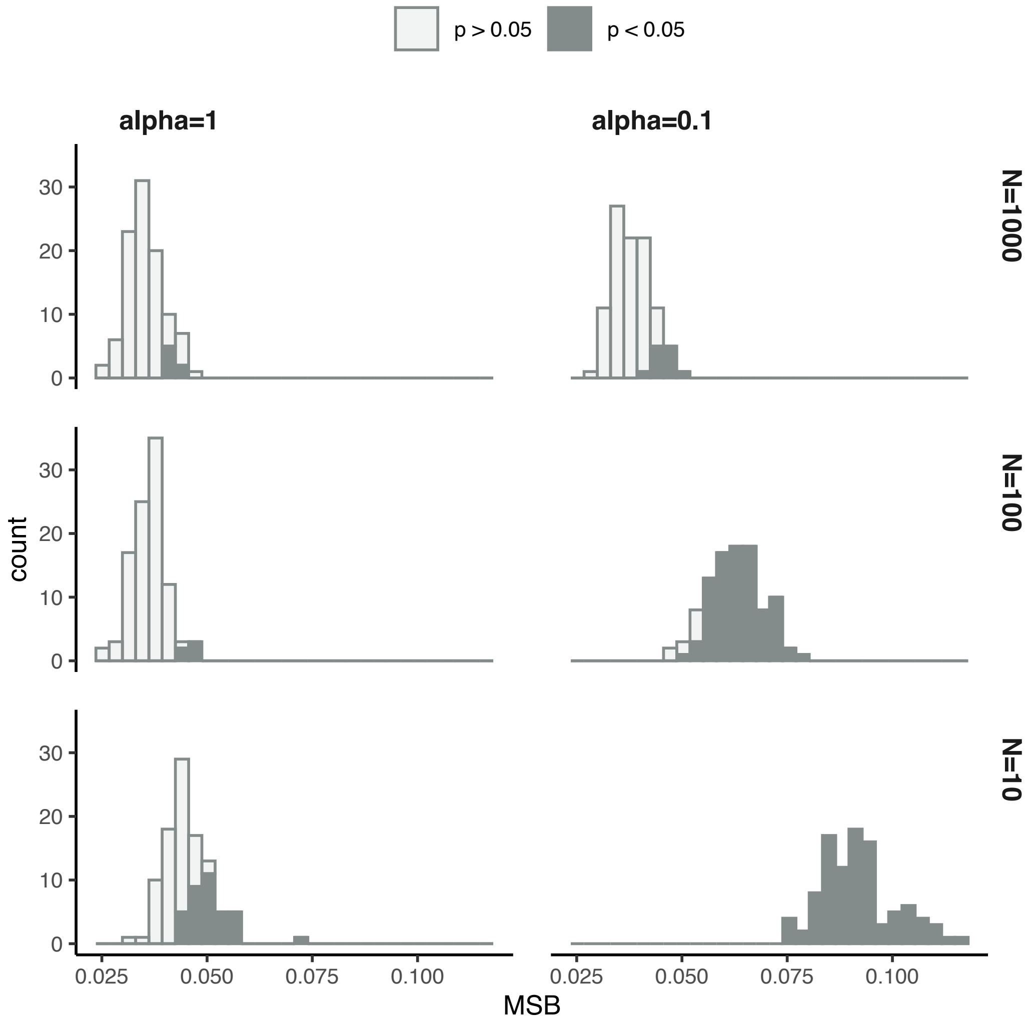


**Supplementary Figure 4. Mutational spectrum bias in simulations.**
Simulations were performed as in the ABC framework, using the neutral mutational spectrum from PAAD and the trinucleotide context composition of *TP53* (see Methods). Each simulation assumed a fixed scaling coefficient $s = 10^6$ and varied the number of driver mutations $N$ and the concentration parameter $a$, which controls the variance of fitness effects across driver mutations. Higher $a$ values correspond to more uniform fitness effects, while lower values produce a more skewed distribution. The total number of possible missense mutations in *TP53*, of which $N$ driver mutations were chosen, equals 2,580. The histograms show the resulting distribution of mutational spectrum bias across 100 simulations per each parameter combination. Filled bars indicate parameter sets where the deviation from the neutral spectrum was significant (likelihood-ratio rest $p < 0.05$ after multiple testing correction).

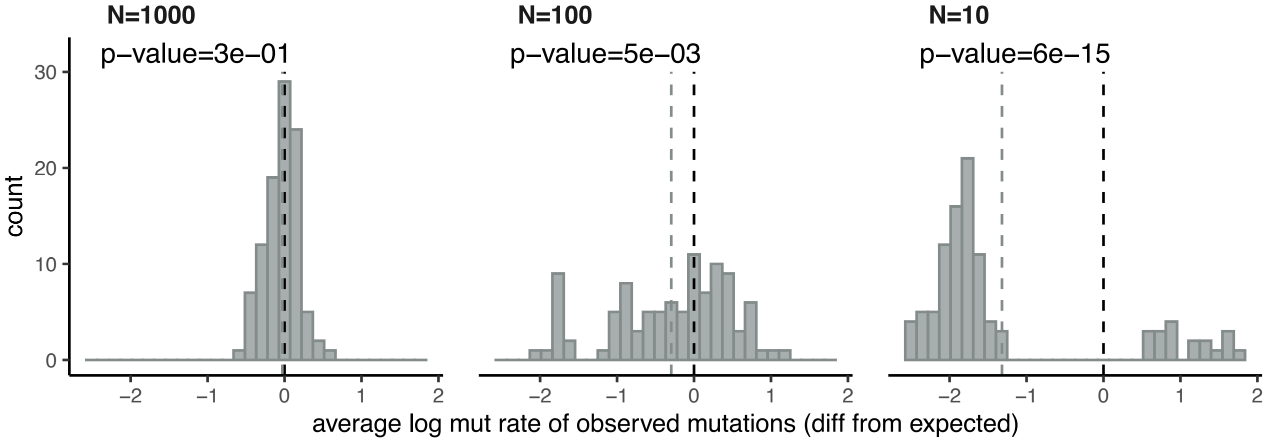


**Supplementary Figure 5. Small number of selection targets shifts the mutational spectrum toward low-mutable contexts.**
Simulations were performed using the PAAD neutral mutational spectrum and the context composition of the *TP53* gene, with fixed parameters $a=1$ and $s=10^6$, and varying the number of driver mutations $N$ (out of 2,580 possible missense mutations). The histograms show the shift in mutability of observed mutations as compared to neutral expectation (see Methods) across 100 simulations for each $N$ value. For each panel, the gray dashed line indicates the mean shift, and p-value (Wilcoxon test) shows deviation from zero.
As $N$ decreases, meaning a smaller target for selection, the spectrum of observed mutations becomes enriched in low-mutability contexts. For very small target sizes ($N=10$), the distribution becomes bimodal: while most simulations exhibit a shift toward low-mutability, occasional cases occur where a highly mutable context is under strong positive selection, further increasing its recurrence.

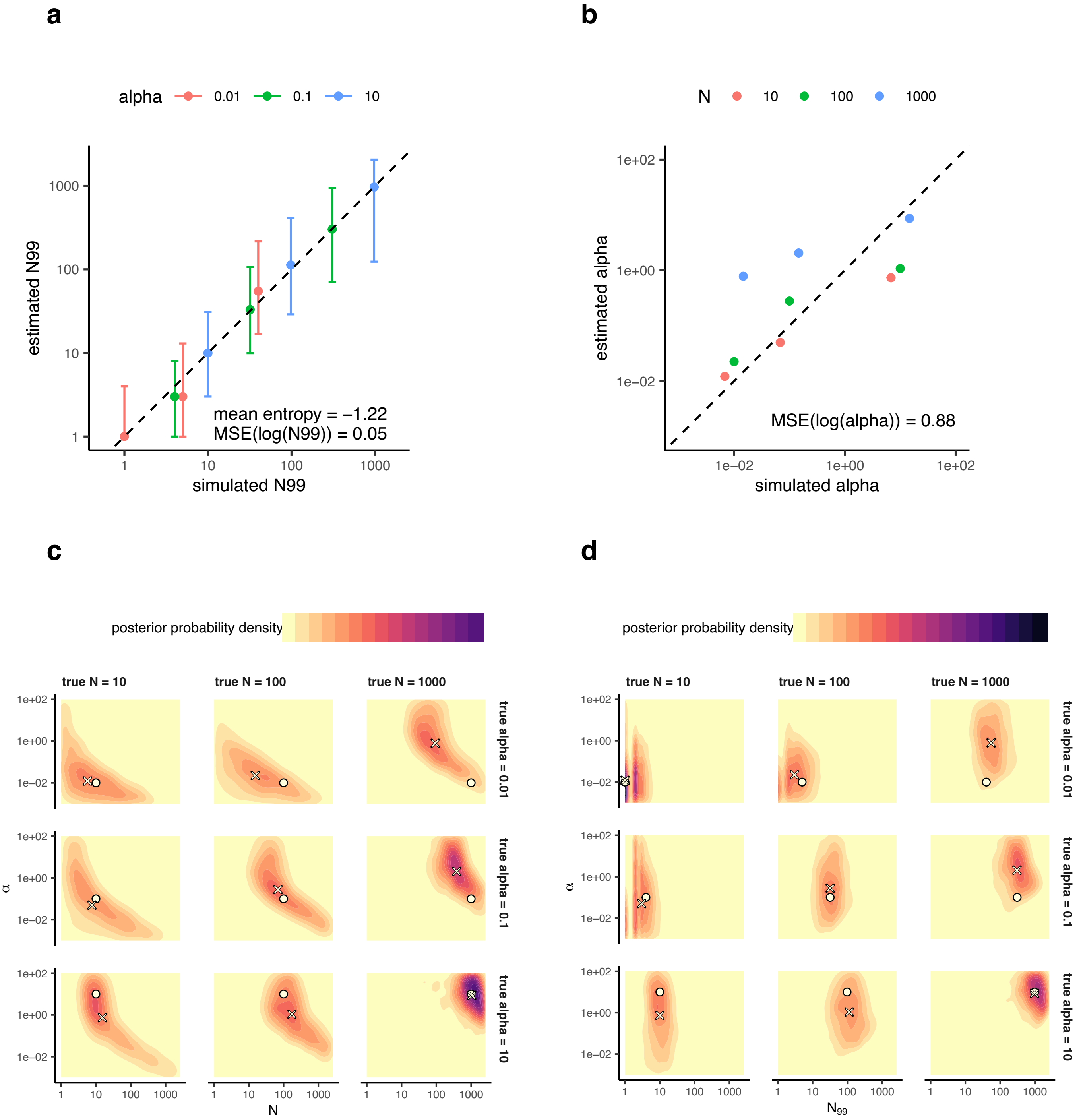


**Supplementary Figure 6. Cross-validation of ABC with four summary statistics (including MSB).**
Simulations were performed using the PAAD neutral mutational spectrum and the trinucleotide context composition of *TP53*. For each combination of $N$ and $a$, the ABC pipeline was run to assess inference accuracy. **a–b**, Correlation between simulated (true) and inferred values for the effective number of driver mutations $N_{99}$ **(a)** and $a$ **(b)**. Errorbars in panel **a** represent the 95% posterior credible intervals for $N_{99}$**. c–d**, Joint posterior distributions of $N$ and $a$ **(c)** and $N_{99}$ and $a$ **(d)**. Circles show the true simulated values; crosses indicate posterior means from ABC inference.

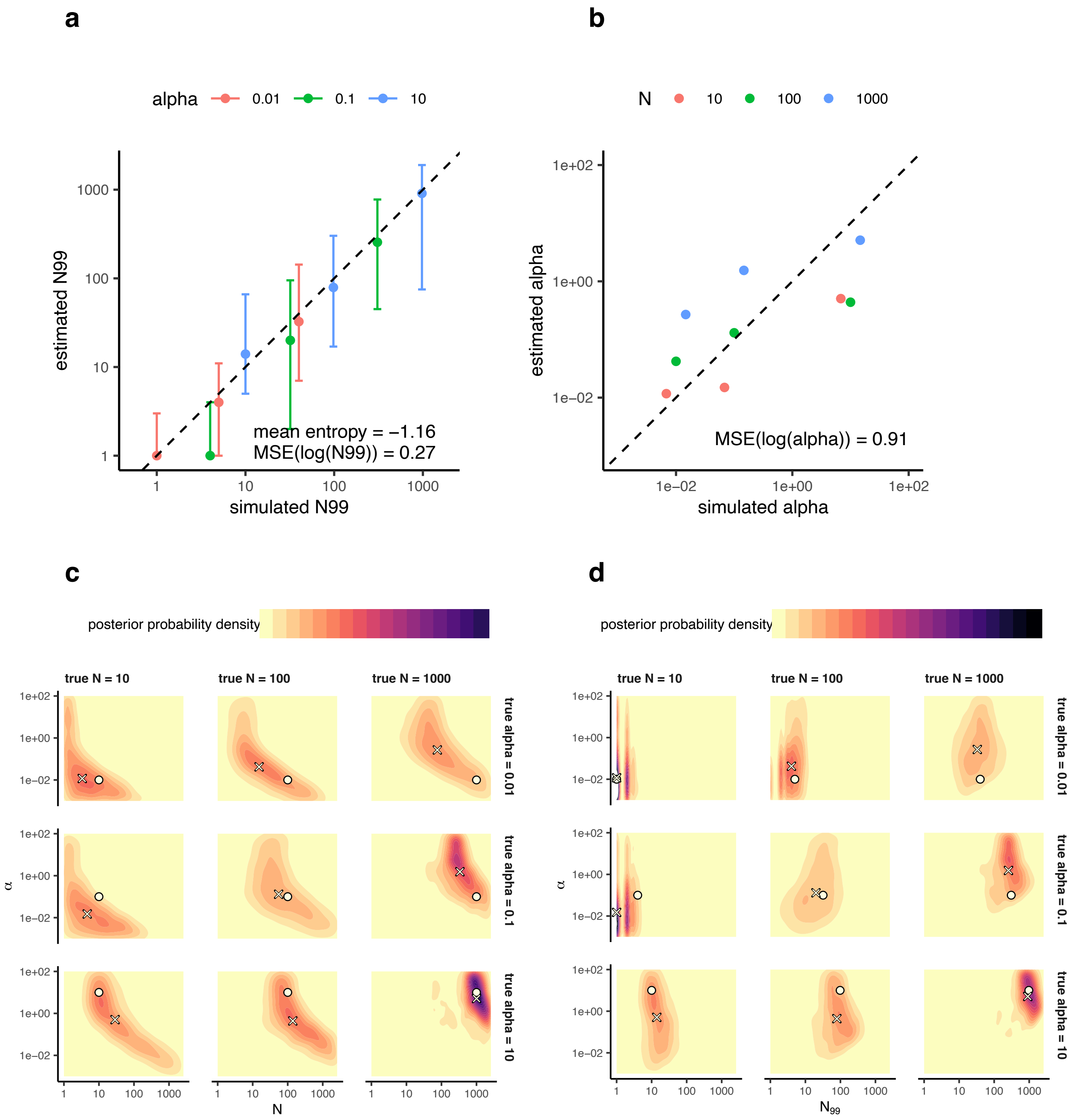


**Supplementary Figure 7. Cross-validation of ABC with three summary statistics (excluding mutational spectrum bias).**
a–b, Correlation between simulated and inferred values for the effective number of driver mutations $N_{99}$ (a) and $\alpha$ (b). Errorbars in panel a represent the 95% posterior credible intervals for $N_{99}$. c–d, Joint posterior distributions of $N$ and $\alpha$ (c) and $N_{99}$ and $\alpha$ (d). Circles show the true simulated values; crosses indicate posterior means from ABC inference.

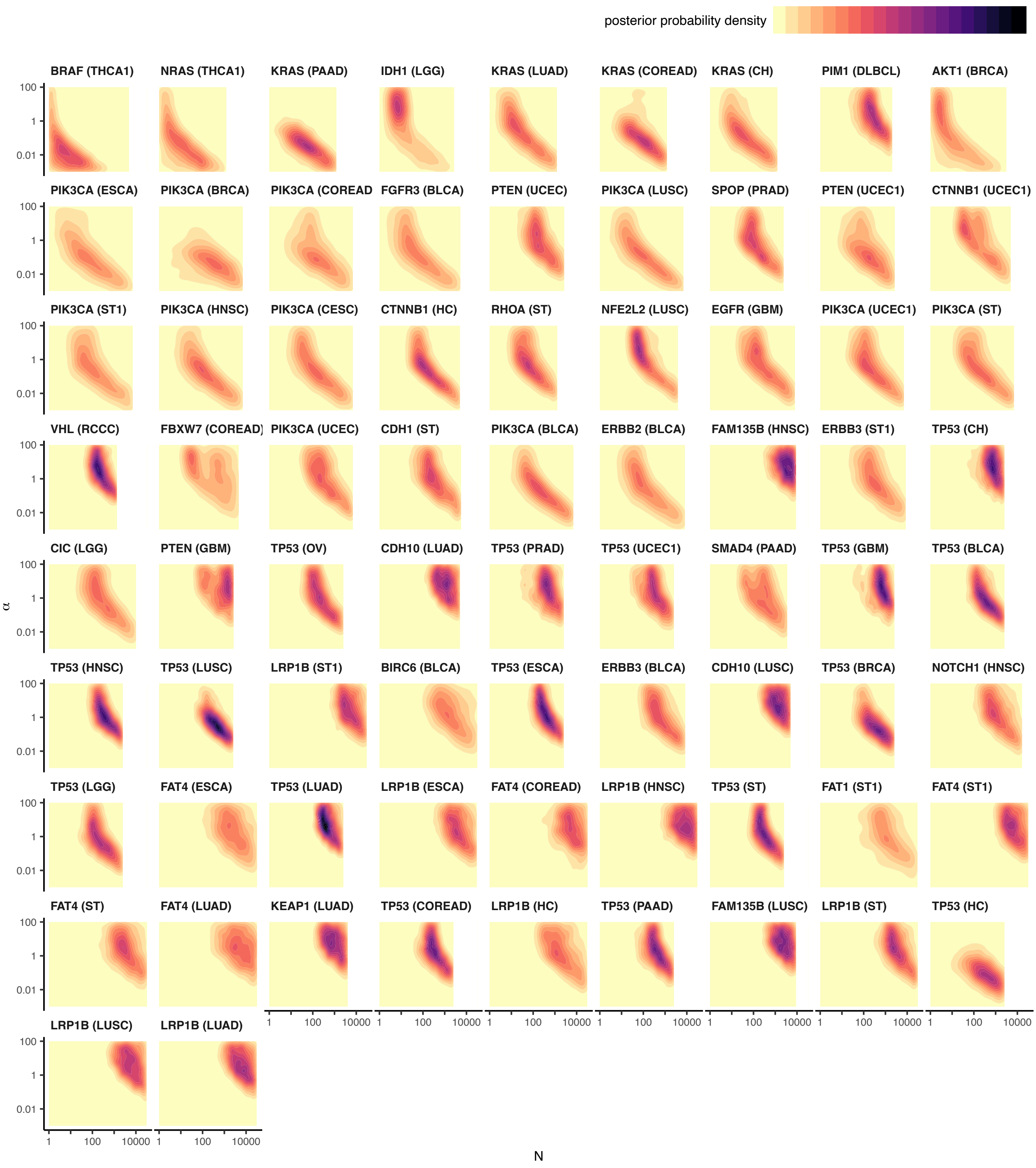


**Supplementary Figure 8. Joint posterior probability distributions of *N* and α in driver genes.** Each panel shows the joint posterior distribution of the number of positively selected sites (*N*) and the concentration parameter ($\alpha$) for one gene in a given tumor type Genes are ordered by decreasing strength of mutational spectrum bias. Only genes with at least 50 observed mutations in a given tumor type are included.

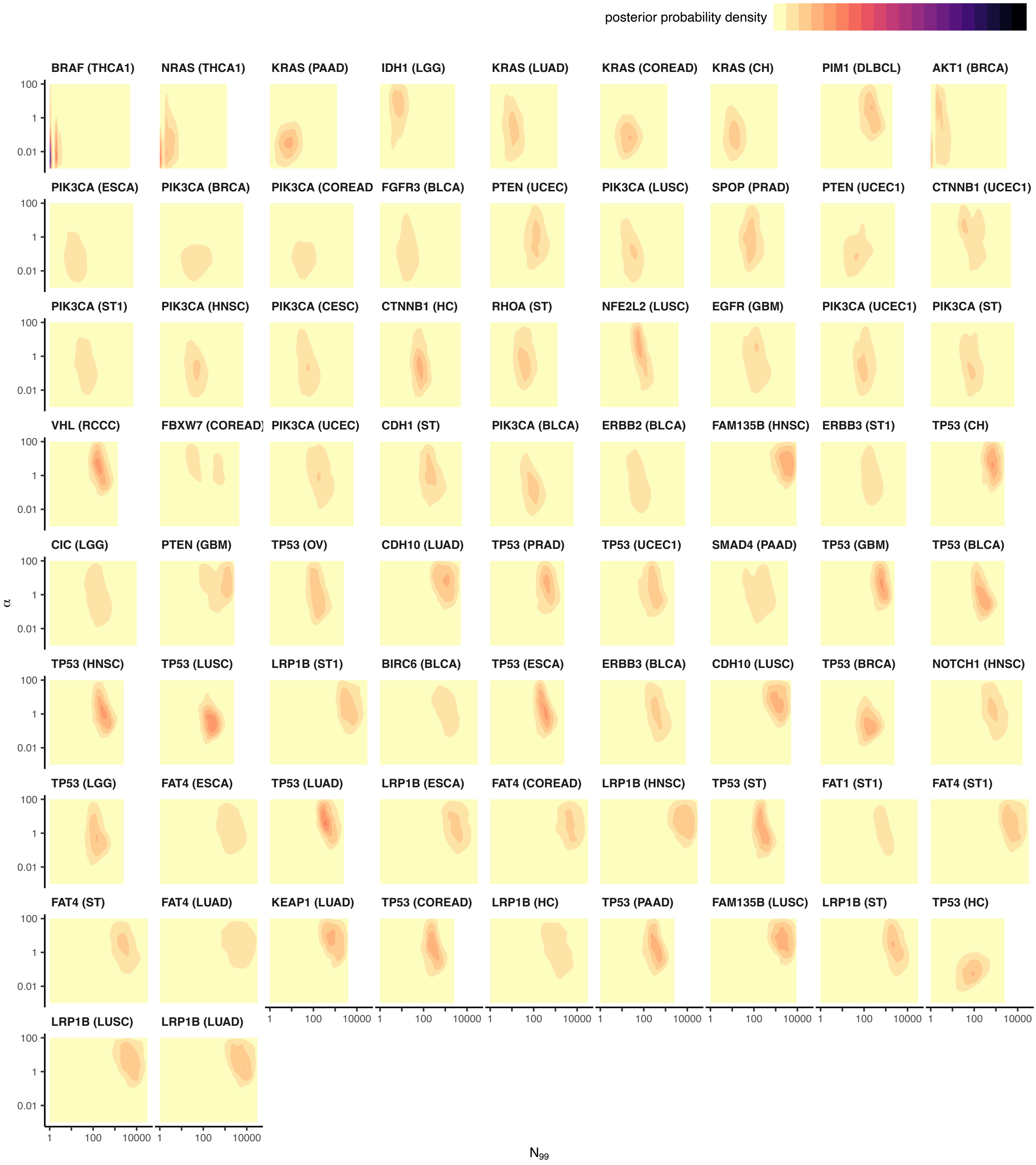


**Supplementary Figure 9. Joint posterior probability distributions of $N_{99}$ and α in driver genes.** Each panel shows the joint posterior distribution of the effective target size ($N_{99}$) and the concentration parameter ($\alpha$) for one gene in a given tumor type Genes are ordered by decreasing strength of mutational spectrum bias. Only genes with at least 50 observed mutations in a given tumor type are included.

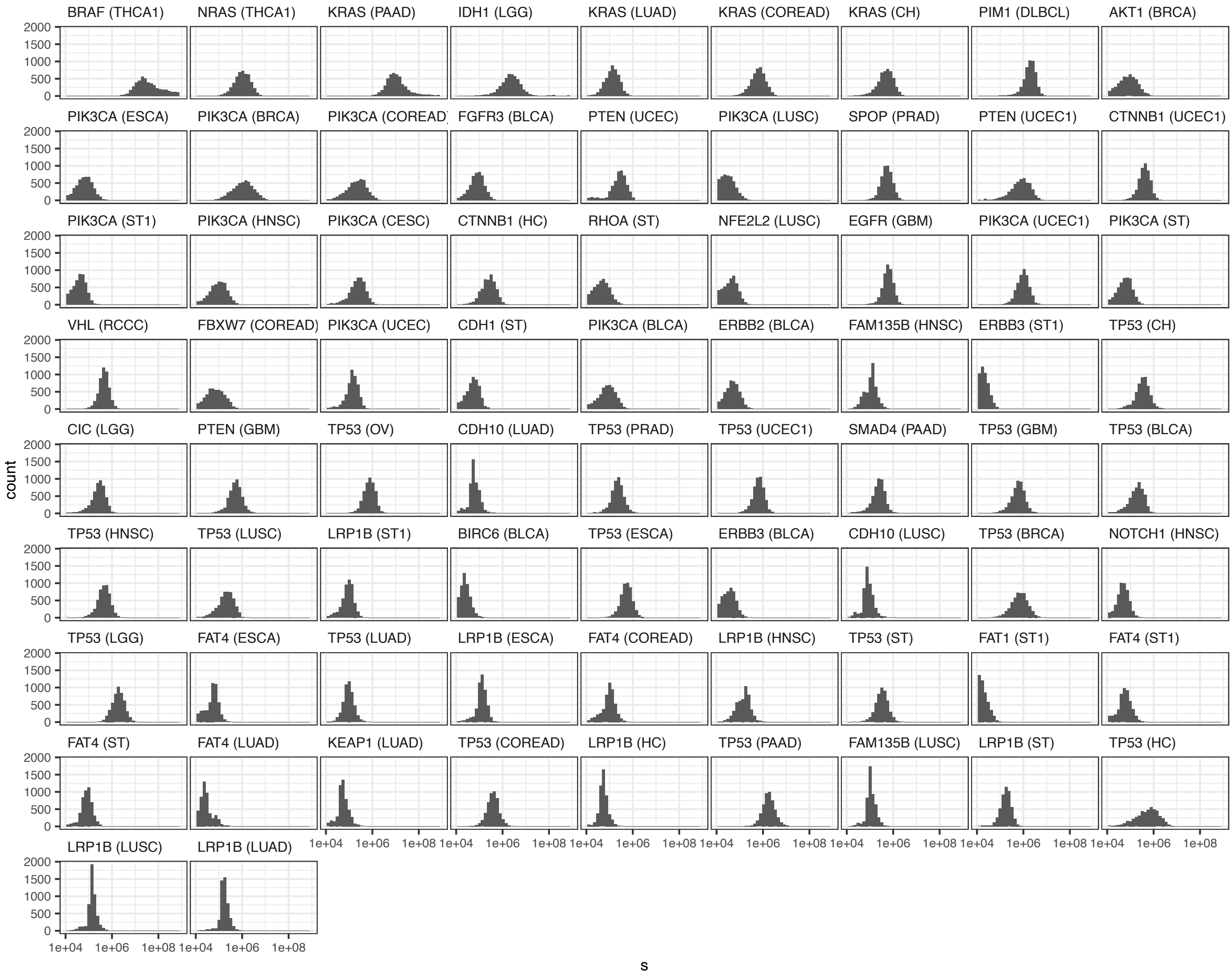


**Supplementary Figure 10. Posterior probability distributions of the scaling coefficient (s) in driver genes.** The genes are sorted by the total number of observed mutations in a given tumor type.

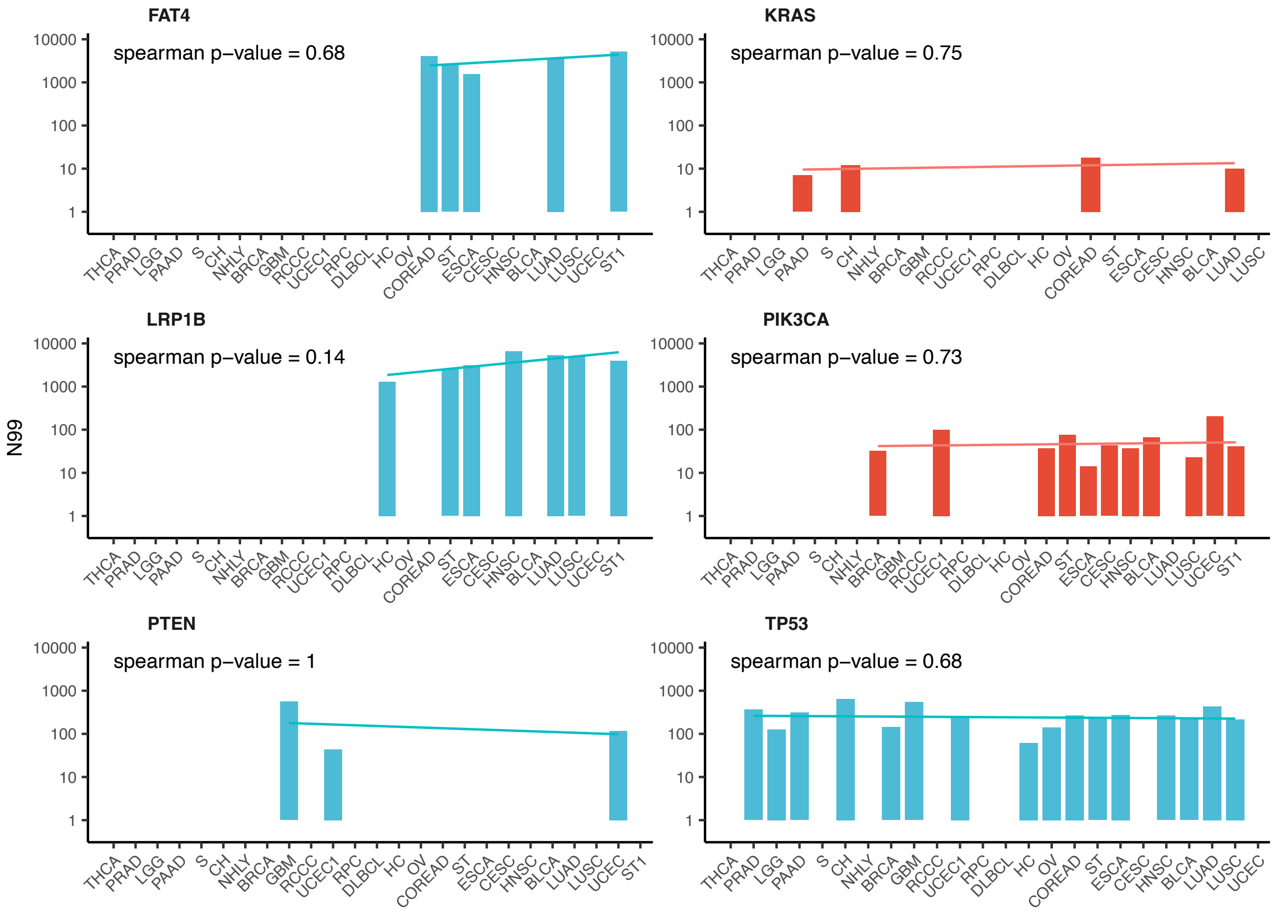


**Supplementary Figure 11. Estimated effective target size ($N_{99}$) in genes annotated as drivers in multiple tumor types.**

Tumor suppressor genes (*FAT4*, *LRP1B*, *PTEN*, and *TP53*) are shown in blue; oncogenes (*KRAS* and *PIK3CA*) are shown in red. Each bar represents the median posterior estimate of $N_{99}$ for a given gene-tumor pair, summarizing the effective number of driver mutations under positive selection. Tumor types are sorted by mutability.

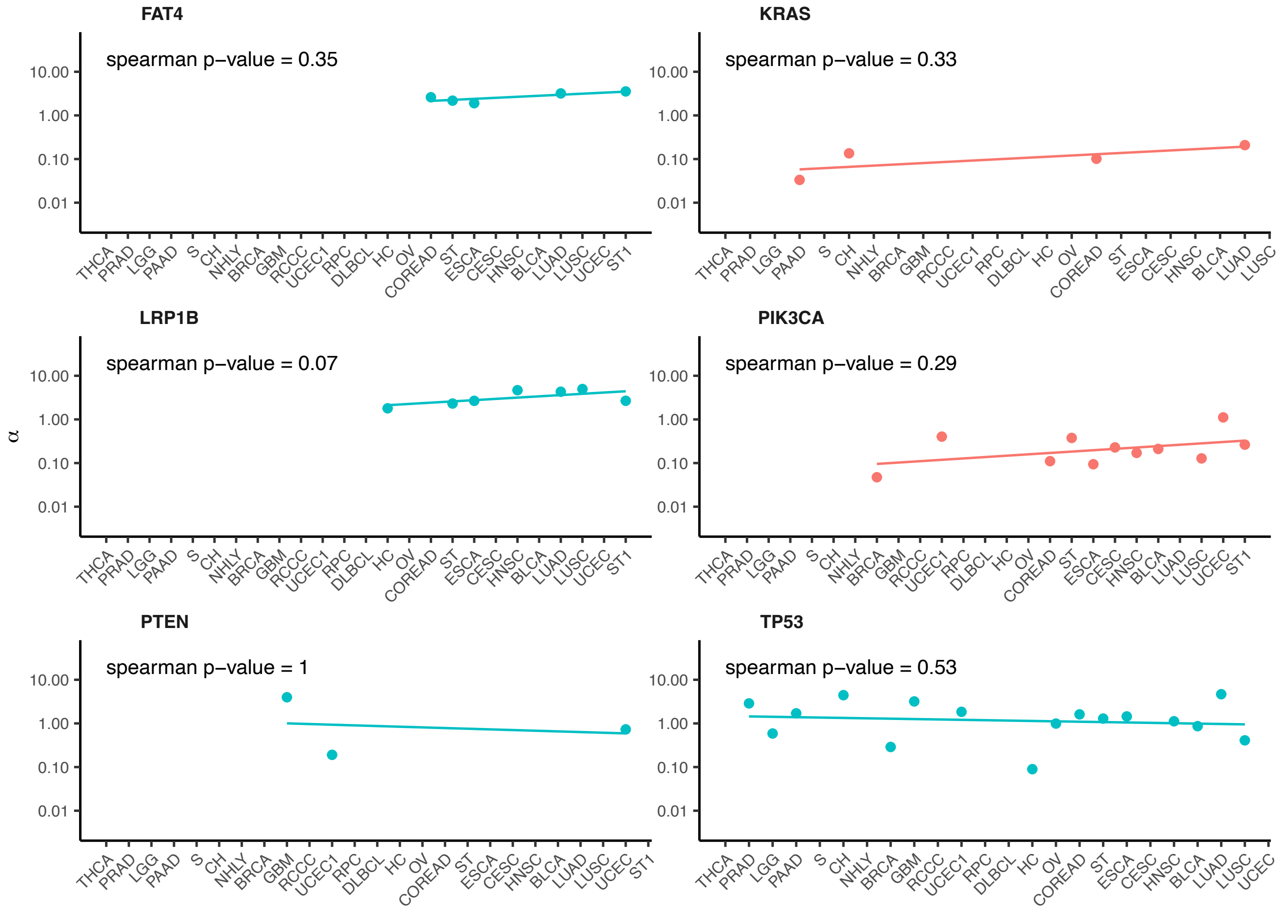


**Supplementary Figure 12. Estimated concentration parameter (α) in genes annotated as drivers in multiple tumor types.**

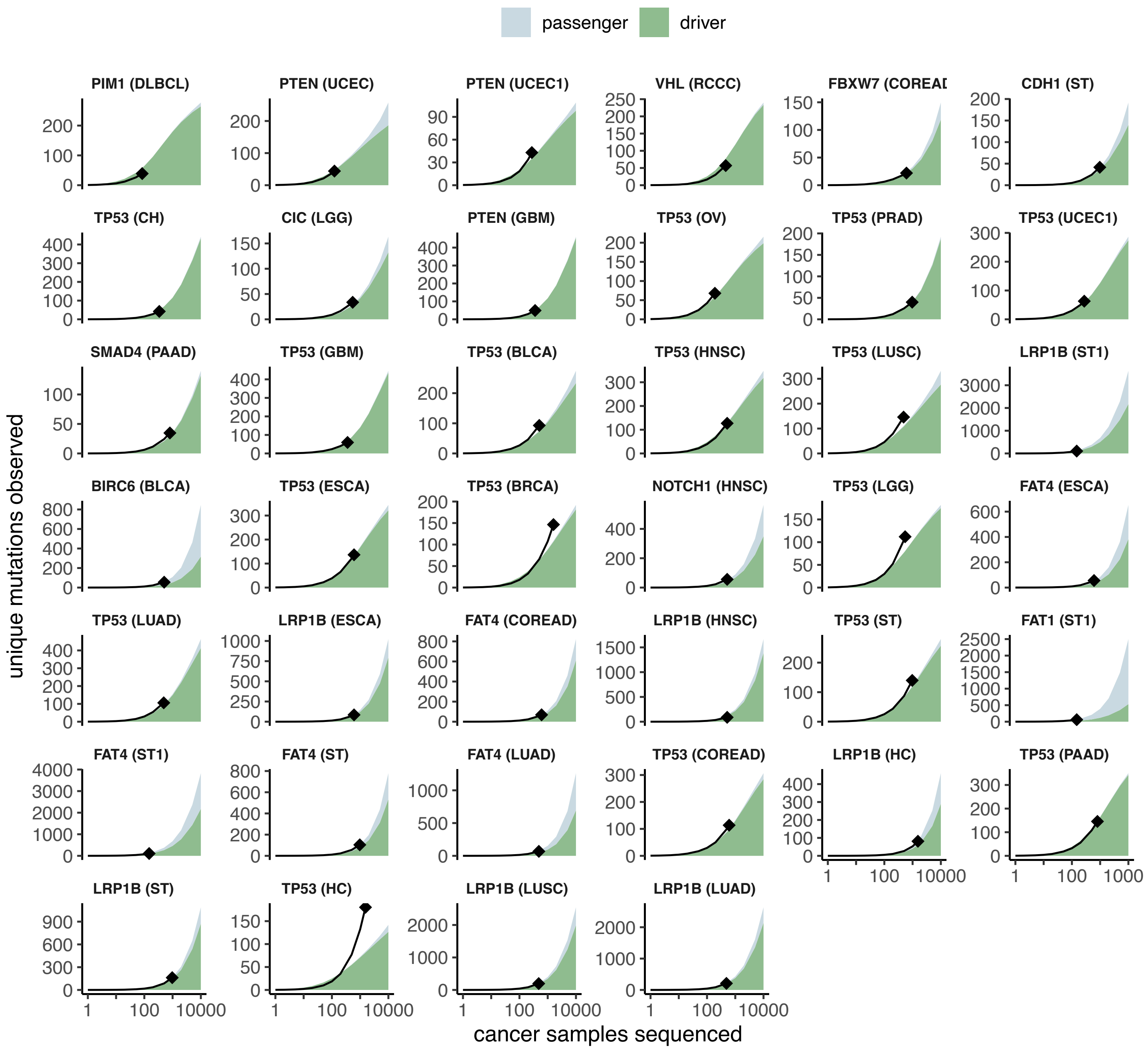


**Supplementary Figure 13. Predicted accumulation of driver mutations with increasing sample size in tumor suppressor genes.**
Diamonds represent the number of unique mutations observed in the original dataset. Black lines indicate rarefaction curves from 1,000 bootstrap resamples of available tumor samples. Green regions show the predicted number of unique driver mutations; gray regions show passenger mutations, based on the average from 5,000 simulations accepted in the last iteration of ABC. Genes are sorted by by decreasing strength of mutational spectrum bias.

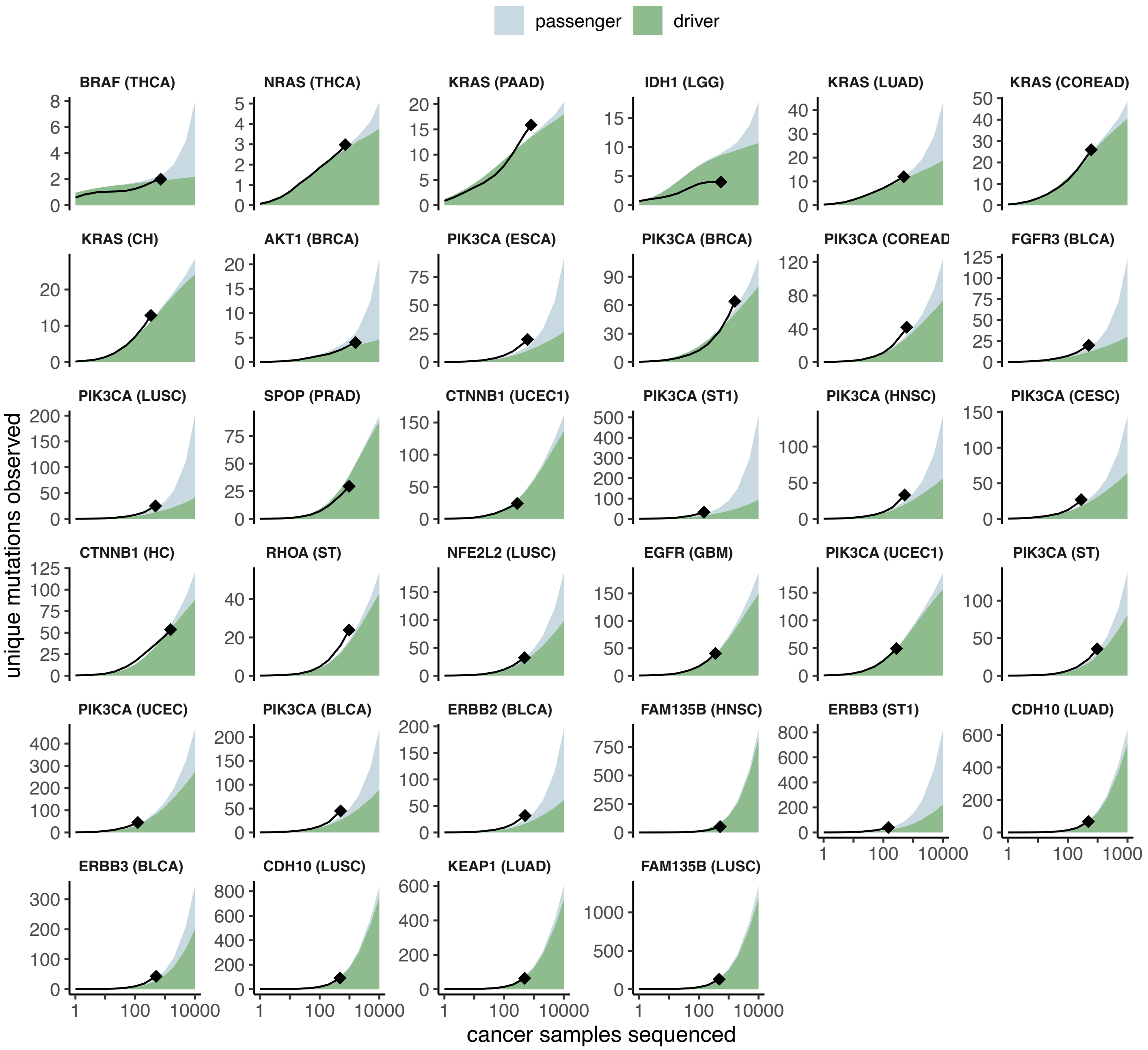


**Supplementary Figure 14. Predicted accumulation of driver mutations with increasing sample size in oncogenes.**
Diamonds represent the number of unique mutations observed in the original dataset. Black lines indicate rarefaction curves from 1,000 bootstrap resamples of available tumor samples. Green regions show the predicted number of unique driver mutations; gray regions show passenger mutations, based on the average from 5,000 simulations accepted in the last iteration of ABC. Genes are sorted by by decreasing strength of mutational spectrum bias.

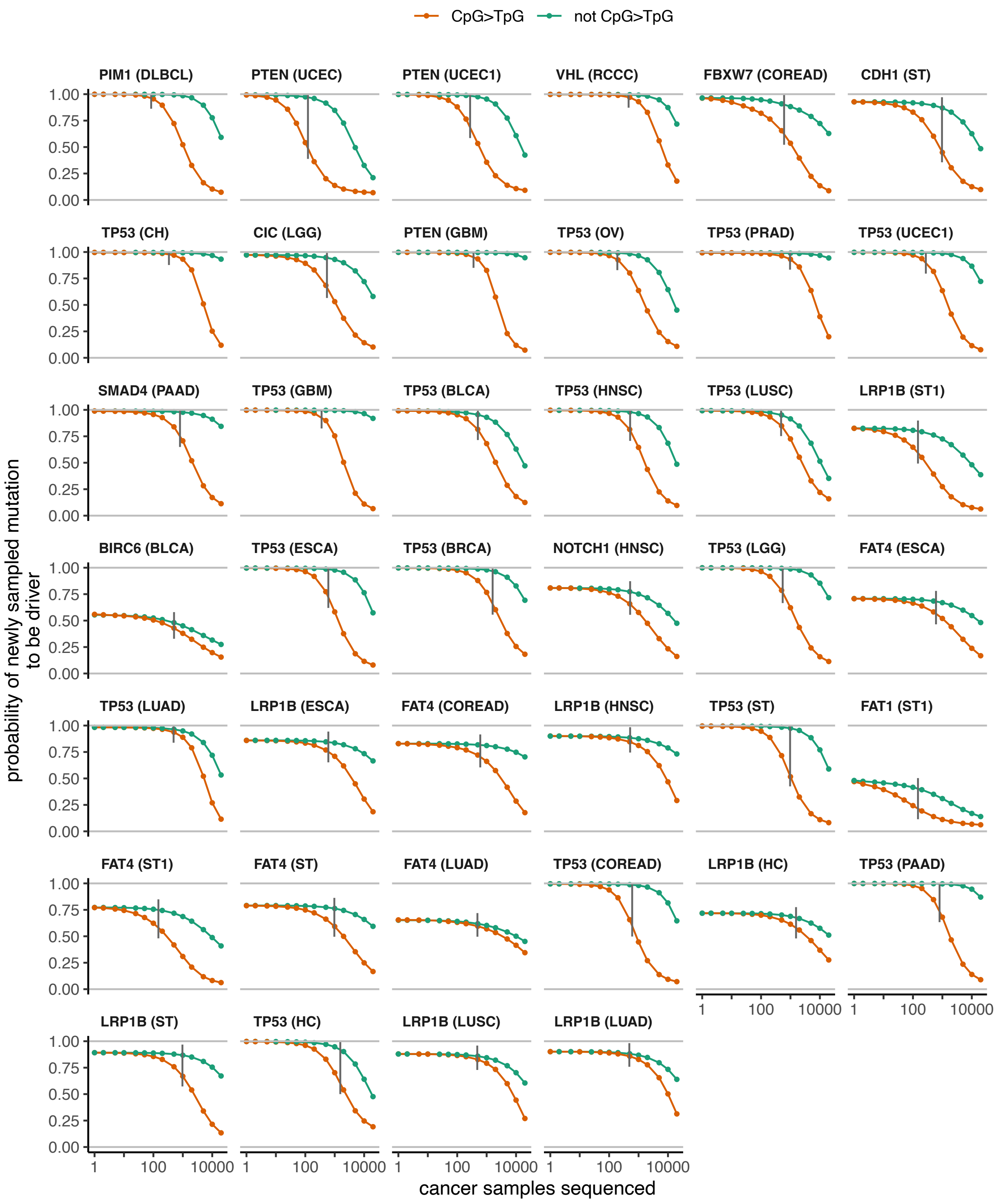


**Supplementary Figure 15. Predicted conditional probabilities for a newly sampled mutation to be driver in tumor suppressor genes.**
For each sample size, the probability that a sampled mutation is under positive selection, given that it wasn't observed in smaller sample sizes. Probabilities were estimated from DFE in driver genes, estimated by ABC (see Methods). Probabilities were calculated separately for C>T in CpG sites, which have high mutation rates in most tumor types, and all other mutation types. Vertical dashed lines indicate the total number of samples available in the analyzed dataset. Genes are sorted by the decreasing strength of mutational spectrum bias.

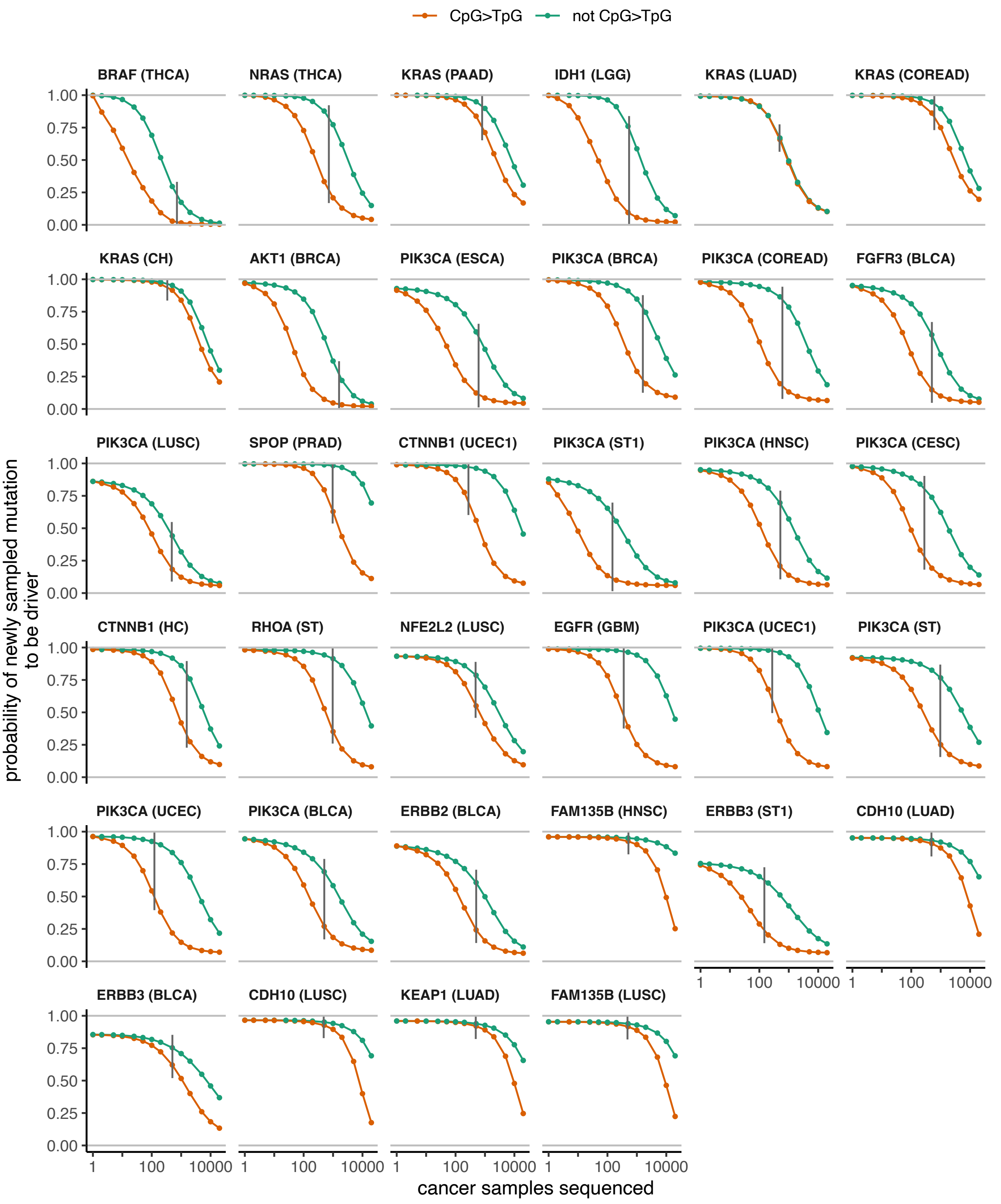


**Supplementary Figure 16. Predicted conditional probabilities for a newly sampled mutation to be driver in oncogenes.**
For each sample size, the probability that a sampled mutation is under positive selection, given that it wasn't observed in smaller sample sizes. For details, refer to the caption of Supplementary Fig. 15.

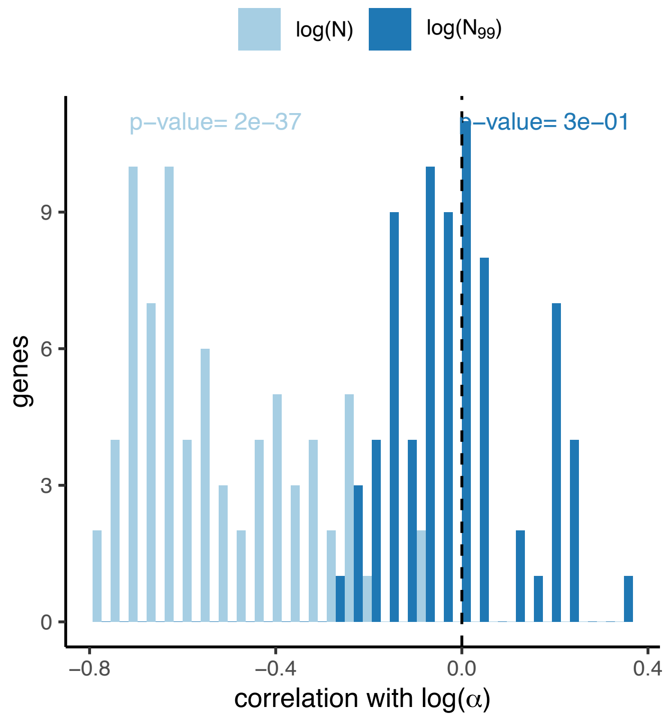


**Supplementary Figure 17. Correlation of $N$ and $N_{99}$ with α in the joint posterior distributions.**

Each point represents one gene in one tumor type. While estimates of $N$ and $\alpha$ are strongly correlated (joint posterior distributions for individual genes are shown in Supplementary Fig. 8), the effective number of driver sites, $N_{99}$, does not depend on α (joint posterior distributions shown in Supplementary Fig. 9).
